# Chromosome-scale scaffolding of the fungus gnat genome (Diptera: *Bradysia coprophila*)

**DOI:** 10.1101/2022.11.03.515061

**Authors:** John M. Urban, Susan A. Gerbi, Allan C. Spradling

## Abstract

**Background:** The fungus gnat, *Bradysia* (*Sciara*) *coprophila*, has compelling chromosome biology. Paternal chromosomes are eliminated during spermatogenesis whereas both maternal X sister chromatids are retained. Embryos start with three copies of the X chromosome, but 1-2 copies are eliminated from somatic cells as part of sex determination, and one is eliminated in the germline to restore diploidy. These developmentally normal events present opportunities to study chromosome movements that are unusual in other systems. To support such studies, we previously generated a highly contiguous optical-map-scaffolded long-read assembly (Bcop_v1) of the male somatic genome. However, the scaffolds were not chromosome-scale, the majority of the assembly lacked chromosome assignments, and the order and orientation of the contigs along chromosomes remained unknown.

**Findings:** Male pupae Hi-C data was used to correct, order, and orient the contigs from Bcop_v1 into chromosome-scale scaffolds, producing the updated assembly, Bcop_v2. Several orthogonal analyses allowed us to (i) identify the corresponding chromosome for each scaffold, (ii) orient them with respect to polytene maps, and (iii) determine that they were highly concordant with the chromosomes they represent. Gene annotations produced for Bcop_v1 were lifted over to Bcop_v2. Chromosomal repeat distributions highlight a potential telomeric sequence. Finally, the Hi-C data shed new light on three “fold-back regions” seen to physically interact in images of polytene X chromosomes.

**Conclusions:** Studies of the unusual chromosome movements in *Bradysia coprophila* will benefit from the updated assembly (Bcop_v2) where each somatic chromosome is represented by a single scaffold.

## Introduction

The lower Dipteran dark-winged fungus gnat, *Bradysia* (*Sciara*) *coprophila*, is an important model system for studying chromosome biology due to its dynamic genome. Most notable is its chromosome cycle. Rather than having either a diploid or haploid set of the same chromosomes (X, II, III, IV, and L) in every cell, the chromosome constitution of a cell varies based on whether it is somatic or germline, male or female, and early embryo or late embryo [1,2]. Regarding somatic or germline, there are chromosomes referred to as the L chromosomes that are only found in the germline [1,2]. Regarding male or female, half of all females, but no males, contain a variant of the X chromosome called the X’ (X prime) [1,2]. Regarding embryo age, embryos start out with three copies of the X chromosome, two of which are paternally derived, and with a variable number of the germline-limited L chromosomes, but later embryos have only 1-2 X chromosomes and no L chromosomes in somatic cells. The L chromosomes are eliminated from all somatic nuclei not set aside for the germline in the 5^th^-6^th^ nuclear division [1,2]. The difference in X chromosome copy number arises because embryos from XX mothers are destined to be male and eliminate two paternal X chromosomes from somatic nuclei in the 7^th^-9^th^ nuclear division whereas embryos produced by X’X mothers are fated to be female and eliminate only one paternal X [1,2]. Accordingly, sex is determined by the chromosome constitution of the mother in this species, which lacks a Y chromosome. Later in development, both the male and female germline restore diploidy for the X by eliminating one paternal copy, and also eliminate one or more L chromosomes to prevent their accumulation [1,2]. Thereafter, the meiotic events in oogenesis appear to proceed normally whereas those in spermatogenesis do not. In meiosis I of spermatogenesis, all paternal somatic chromosomes are eliminated in a bud of cytoplasm while the L chromosomes appear to escape this imprinting effect and are retained with the maternal set of somatic chromosomes at the single pole of a naturally occurring monopolar spindle [1,2]. In meiosis II, both maternally-derived sister chromatids of the X chromosome undergo developmentally programmed nondisjunction and are retained at one pole of a bipolar spindle [1,2]. Rather than four products, these unusual events lead to one product of male meiosis: a sperm that contains only maternally derived somatic chromosomes (X, II, III, and IV), that is diploid for X, haploid for the autosomes (II, III, and IV), and variable for L chromosomes. In addition to the unusual chromosome cycle, *Bradysia coprophila* larval salivary glands have highly polytene chromosomes with over 8000 copies in each nucleus [3], and developmentally-programmed gene amplification [4,5]. Finally, there is mounting evidence that this genome may be a great model for studying horizontal gene transfer (HGT) in insects [6–8]. Overall, fungus gnats present many opportunities to study chromosome biology across development and evolution.

Previously, we published a highly contiguous assembly of the somatic chromosomes (X, II, III, and IV) named Bcop_v1 [9]. We targeted the somatic genome in males as it presented the lowest complexity version of the genome, lacking both the L and X’ chromosomes, which have since been the focus of other studies [6,10]. Specifically, we sequenced genomic DNA from male embryos using long-read (PacBio RS II SMRT and Oxford Nanopore Technologies MinION) and short-read (Illumina) technologies [9]. As part of that process, we assembled the sequencing data ∼100 different ways, and used many evaluations to determine the best assembly and polishing pipelines for our datasets [9]. Two assemblies with the best evaluation scores were scaffolded with optical maps from BioNano Genomics, then subject to further evaluations[9]. The BioNano-scaffolded assembly with the best evaluation scores, Bcop_v1 (NCBI GenBank GCA_014529535.1, WGS VSDI01), had a contig NG50 of ∼2.4 Mb and scaffold NG50 of ∼8.2 Mb (Table 1) [9]. Since, in *Bradysia coprophila*, late male embryos have a single copy of the X, but are diploid for the autosomes, it was possible to classify each contig as either X-linked or autosomal using sequencing depth [9]. Moreover, using the known chromosomal locations of a handful of sequences, we were able to anchor ∼28-33% of the autosomal sequences into chromosomes II, III, and IV [9]. Overall, nearly 50% of the expected genome size was anchored into chromosomes based on read depth and known sequence locations [9]. Nevertheless, chromosome assignments are not available for the majority of that assembly (Bcop_v1). Moreover, Bcop_v1 lacks any information regarding the ordering and orientation of the contigs along chromosome sequences. To better facilitate studies into gene amplification and the unusual chromosome dynamics in *Bradysia coprophila*, our goal was to develop better models of the somatic chromosome sequences. We approached this by using chromosome conformation capture with deep sequencing (Hi-C) [11] to scaffold the chromosomes as has been successfully done by many others [12–14]. Specifically, Hi-C is powerful at putting chromosome sequences together from a collection of sub-chromosomal sequences because of three major attributes of *in vivo* chromosome interactions captured by Hi-C data: (i) the interactions arbitrarily yield Mb-scale and chromosome-scale information without the need for sequencing ultra-long DNA molecules, (ii) intra-chromosomal interactions are much more frequent than inter-chromosomal interactions, making it possible to use interaction frequencies to cluster contigs into ‘chromosome groups’, and (iii) interaction frequencies decay with distance, making it possible to use them to order and orient the contigs in a chromosome group in the order and orientations they most likely appear along a chromosome [11–14].

**Table 1:**
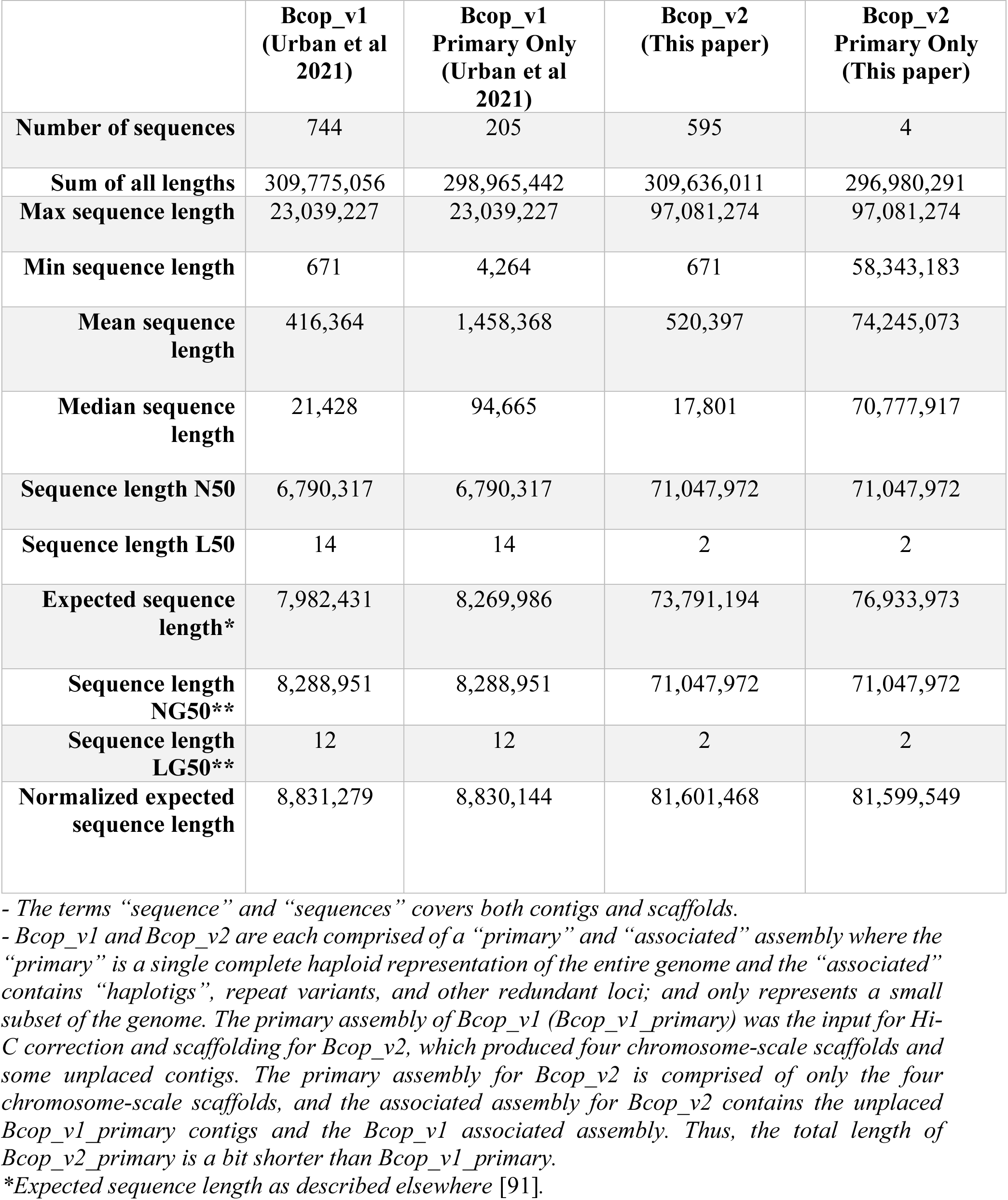
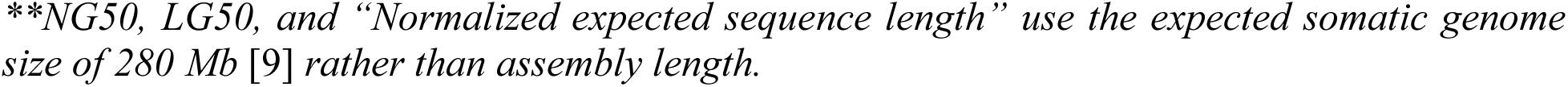
Contiguity Statistics comparing the assembly before (Bcop_v1) and after (Bcop_v2) Hi-C scaffolding (Bcop_v2).

|  | **Bcop_v1**<br>**(Urban et al**<br>**2021)** | **Bcop_v1**<br>**Primary Only**<br>**(Urban et al**<br>**2021)** | **Bcop_v2**<br>**(This paper)** | **Bcop_v2**<br>**Primary Only**<br>**(This paper)** |
| --- | --- | --- | --- | --- |
| **Number of sequences** | 744 | 205 | 595 | 4 |
| **Sum of all lengths** | 309,775,056 | 298,965,442 | 309,636,011 | 296,980,291 |
| **Max sequence length** | 23,039,227 | 23,039,227 | 97,081,274 | 97,081,274 |
| **Min sequence length** | 671 | 4,264 | 671 | 58,343,183 |
| **Mean sequence**<br>**length** | 416,364 | 1,458,368 | 520,397 | 74,245,073 |
| **Median sequence**<br>**length** | 21,428 | 94,665 | 17,801 | 70,777,917 |
| **Sequence length N50** | 6,790,317 | 6,790,317 | 71,047,972 | 71,047,972 |
| **Sequence length L50** | 14 | 14 | 2 | 2 |
| **Expected sequence**<br>**length\*** | 7,982,431 | 8,269,986 | 73,791,194 | 76,933,973 |
| **Sequence length**<br>**NG50\*\*** | 8,288,951 | 8,288,951 | 71,047,972 | 71,047,972 |
| **Sequence length**<br>**LG50\*\*** | 12 | 12 | 2 | 2 |
| **Normalized expected**<br>**sequence length** | 8,831,279 | 8,830,144 | 81,601,468 | 81,599,549 |

- The terms “sequence” and “sequences” covers both contigs and scaffolds.
- Bcop\_v1 and Bcop\_v2 are each comprised of a “primary” and “associated” assembly where the “primary” is a single complete haploid representation of the entire genome and the “associated” contains “haplotigs”, repeat variants, and other redundant loci; and only represents a small subset of the genome. The primary assembly of Bcop\_v1 (Bcop\_v1\_primary) was the input for Hi-C correction and scaffolding for Bcop\_v2, which produced four chromosome-scale scaffolds and some unplaced contigs. The primary assembly for Bcop\_v2 is comprised of only the four chromosome-scale scaffolds, and the associated assembly for Bcop\_v2 contains the unplaced Bcop\_v1\_primary contigs and the Bcop\_v1 associated assembly. Thus, the total length of Bcop\_v2\_primary is a bit shorter than Bcop\_v1\_primary.
\*Expected sequence length as described elsewhere [91].
*\*\*NG50, LG50, and “Normalized expected sequence length” use the expected somatic genome* *size of 280 Mb [9] rather than assembly length.*

Overall, we present here the first chromosome-scale genome assembly for *Bradysia coprophila*, specifically reporting the first full-length scaffolds for the somatic chromosomes (X, II, III, IV). In addition, we lifted over the annotations from the former genome assembly, allowing studies in progress by multiple groups to seamlessly transition to the updated genome assembly. Through several approaches, we demonstrate the chromosome identity and high structural accuracy of the chromosome-scale scaffolds, including through inspecting Hi-C interaction frequencies, correlation with polytene maps, how repeats are distributed, read depth, evolutionary expectations, and special features of the X chromosome. The termini of the chromosome-scale scaffolds hint that telomeres may be composed of Long Complex Terminal Tandem Repeat (LCTTR) sequences as seen in the non-biting midge *Chironomus* and the mosquito *Anopheles gambiae* [15,16], but also arrays of retrotransposon-related sequences. In addition, the Hi-C maps show three loci, separated by >10 Mb each, with high frequency interactions *in vivo*, potentially illuminating the loci of the three “fold-back regions” seen to physically interact in images of the polytenized X chromosome. Finally, multiple genomic datasets demonstrate that recently-discovered “alien” genes [7] are genuine parts of the chromosome-scale scaffolds, not contamination or mis-assemblies. The Hi-C data, the chromosome-scale genome sequence, its gene annotations, and insights produced here will be immediately useful to the growing research community interested in the unique biology of *Bradysia coprophila* as well as the broader research community interested in Dipteran evolution and comparative genomics.

## Results

### Hi-C guided assembly correction and chromosome-scale scaffolding of the reference genome

To update the previous reference genome, Bcop_v1 [9] (for background on Bcop_v1, see the Introduction, Table 1), we generated Hi-C data for chromosome-scale scaffolding. Approximately 115.2 million 101bp paired-end reads were produced from the chimeric DNA molecules obtained with a commercially-available chromosome conformation capture (Hi-C) procedure from Phase Genomics (Proximo Hi-C kit) performed on whole male pupae (see Methods). The paired-end Hi-C reads were first used to produce a corrected assembly (called “Bcop_v1_corrected”) by aligning them to the 205 primary contigs of Bcop_v1 (“Bcop_v1_primary”) [9], then manually inspecting the Hi-C interaction frequency signal within the Bcop_v1 contigs for disruptions in the expected pattern corresponding to putative mis-joined regions (Fig. 1A-C). Bcop_v1 had 22 contigs with 46 such disruptions in the Hi-C signal (examples in Fig. 1B-C). The 22 contigs were “corrected” by breaking them into “sub-contigs” at the putative mis-joined regions. Of the 46 “debris” regions where breaks were made in Bcop_v1 contigs, 32 were adjacent to BioNano optical map gaps, i.e. regions joined together previously by BioNano optical maps in Bcop_v1, with gap lengths ranging from ∼4.4 to ∼674 kb. However, of the 156 BioNano gaps in Bcop_v1, there were 124 ranging from 25 bp to ∼400 kb that were conserved after the correction step, 70 of which were on the same 22 contigs that had “debris” regions and 54 that were elsewhere within Bcop_v1. Thus, the regions joined by BioNano optical maps mostly did not present as putative mis-joins. Lower mapping quality from flanking repeats around BioNano gaps may have simply given rise to spurious Hi-C signals more often than random. In summary, the majority of mis-join signals (∼70%) flanked a minority of assembly gaps, and mis-join signals affected a minority of contigs (10.7%) that were broken up by Hi-C-guided correction. This step transformed the input assembly (Bcop_v1_primary) that had 205 contigs into the corrected assembly (Bcop_v1_corrected), which had 297.

**Figure 1.**
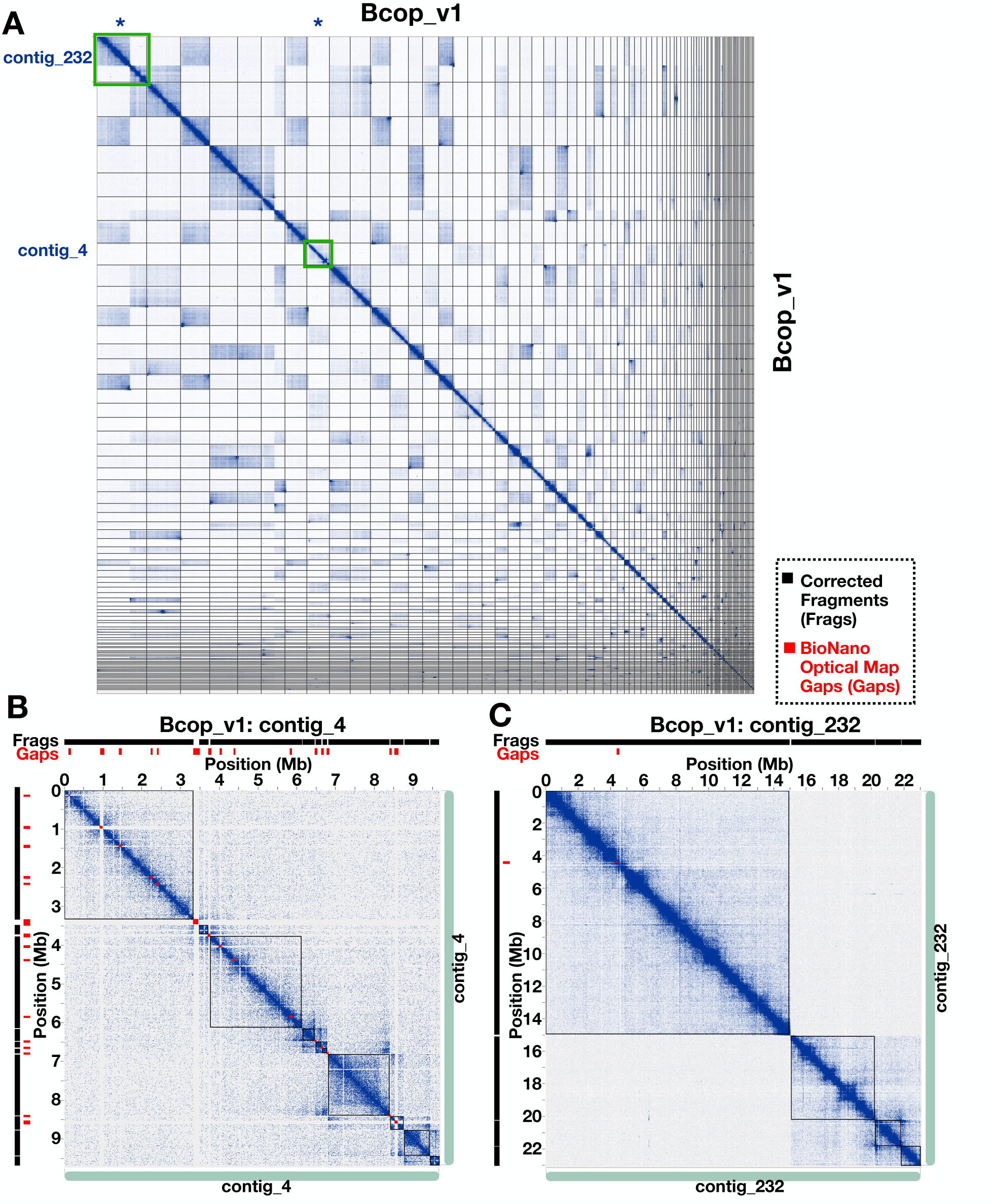
Hi-C guided correction of contigs from Bcop_v1 **(A)** The entire Hi-C map of all primary scaffolds and contigs in Bcop_v1 ordered from longest to shortest before correction and scaffolding. Scaffolds from B and C are highlighted by green boxes. **(B)** Shows example of disrupted Hi-C signals related to joins made by scaffolding with BioNano Genomics optical maps and associated gaps. This is contig_4, a ∼9.7 Mb contig with 14 optical map gaps that was broken into 10 corrected fragments with the 9 breaks correlated with gap locations. New corrected contigs are outlined by black squares and optical map gaps are represented by red squares. Both are also displayed above and to the left of the Hi-C map. The bottom and right show seafoam-colored bars representing the full original Bcop_v1 contig. **(C)** Example of disrupted Hi-C signal that occurred independent of optical map gaps within contiguous sequence. The Hi-C signal forms two obvious sequences that interact within themselves but that are significantly depleted of interactions with each other. The black squares outline the new contigs after correction. The self-interacting sequence on the bottom right has additional Hi-C signals that suggested possible structural errors; hence it being broken up into additional fragments. Fragments, optical map gaps, and the original contig are outlined and represented as in B.

The Phase Genomics’ Proximo Hi-C genome scaffolding platform was then used to create chromosome-scale scaffolds from the 297 contigs in the corrected input assembly (Bcop_v1_corrected; Fig. 2A). After scaffolding and post-processing (see Methods), there were four primary chromosome-scale scaffolds, each spanning 58-71 Mb and another 52 input contigs that summed to ∼1.8 Mb, which were not placed in the chromosome-scale scaffolds (Table 1; Fig. 2B; see Methods). Thus, the vast majority of the ∼299 Mb of primary sequence from Bcop_v1 (Table 1) input into the Hi-C scaffolding process was integrated into the four chromosome-scale scaffolds that summed to ∼297 Mb. In addition, there were 539 “associated contigs” from Bcop_v1 [9], largely comprised of haplotigs and repeat variants, that were not included during the Hi-C scaffolding process (see Methods), which summed to 10.8 Mb and were ∼20 kb on average. As a semi-final step in producing the updated reference genome, Bcop_v2, these 539 contigs were also added back to the assembly. Moving forward, the four chromosome-scale scaffolds are referred to as the “primary assembly” for Bcop_v2 whereas the “associated assembly” for Bcop_v2 is composed of the 52 unplaced input contigs and the 539 “associated contigs” from Bcop_v1, totaling 591 “associated contigs” for Bcop_v2. Most of the associated assembly is comprised of haplotigs and repeat variants. Researchers interested mainly in the four chromosome-scale scaffolds can opt to use or ignore the associated assembly. The four primary chromosome scaffolds make up ∼96% of the entire assembly with lengths of 58.3, 70.5, 71.0, and 97.1 Mb (Tables 1 and 2), and together with the 591 associated contigs have an expected size of ∼81 Mb, and an NG50 length of ∼71 Mb pertaining to an LG50 count of only 2 scaffolds (Table 1). In the updated assembly (Bcop_v2), gaps from the original BioNano scaffolding process (for Bcop_v1) ranged from 25 bp to ∼662 kb with a gap N50 of ∼101.5 kb. Hi-C scaffolding introduced 241 gaps, all arbitrarily set to 100 bp, 5 of which are on a short ∼363 kb scaffold. In total, there are 367 Hi-C and/or optical map gaps across Bcop_v2, 360 of which are within the four chromosome-scale scaffolds. In summary, Hi-C scaffolding transformed the corrected assembly into a preliminary version of Bcop_v2, which had 4 primary scaffolds and 591 associated contigs. The scaffolds were further oriented using the positions of known sequences as described in a subsequent section.

**Figure 2:**
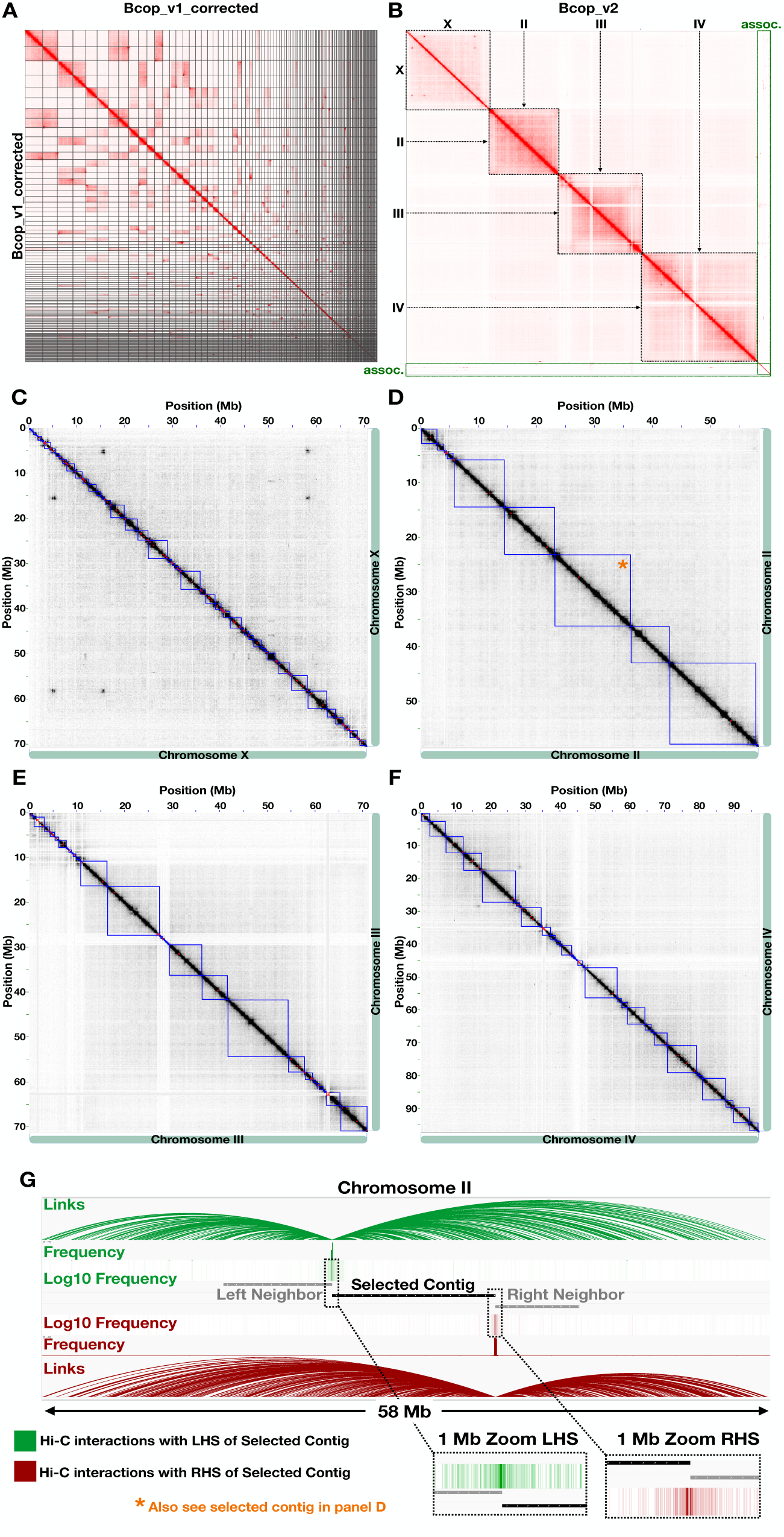
Hi-C signal shows high ‘chromosome specificity’ and ‘structural accuracy’ of the chromosome-scale scaffolds. **(A)** Interaction frequencies are visualized across Bcop_v1_corrected (Bcop_v1 after Hi-C guided correction), which was the input to Hi-C scaffolding. **(B)** Interaction frequencies visualized across the final version of Bcop_v2, after polytene orientation and with associated/unplaced contigs. Squares are drawn around boundaries of the four chromosome-scale scaffolds. **(C, D, E, F)** Interaction frequencies across the X, II, III, and IV chromosome scaffolds, respectively. Blue squares correspond to individual contigs from Bcop_v1_corrected that make up the scaffolds. The asterisk in D corresponds to the selected contig in G. **(G)** An example of the interaction frequencies between the ends of a selected contig and the rest of the chromosome-scale scaffold. The selected contig shown was from chromosome II (see asterisk in panel D). The selected contig is colored black whereas the two neighboring contigs are grey. Links (arcs) tracks correspond to a summary of loci that were linked together by paired-end reads where at least one mate maps to the given contig end. The “Frequency” tracks are bar plots corresponding to the number of times each bin across the chromosome was linked to the given contig end. However, interaction frequency decays exponentially with distance. Thus, the “Log10 Frequency” across the chromosome is also shown as a heatmap, and zoom-ins of the Log10 Frequency heatmaps in 1 Mb regions centered on each contig end are shown for higher resolution. Green and red tracks correspond to interactions and interaction frequencies with the left end and the right end of the selected contig, respectively.

**Table 2:**
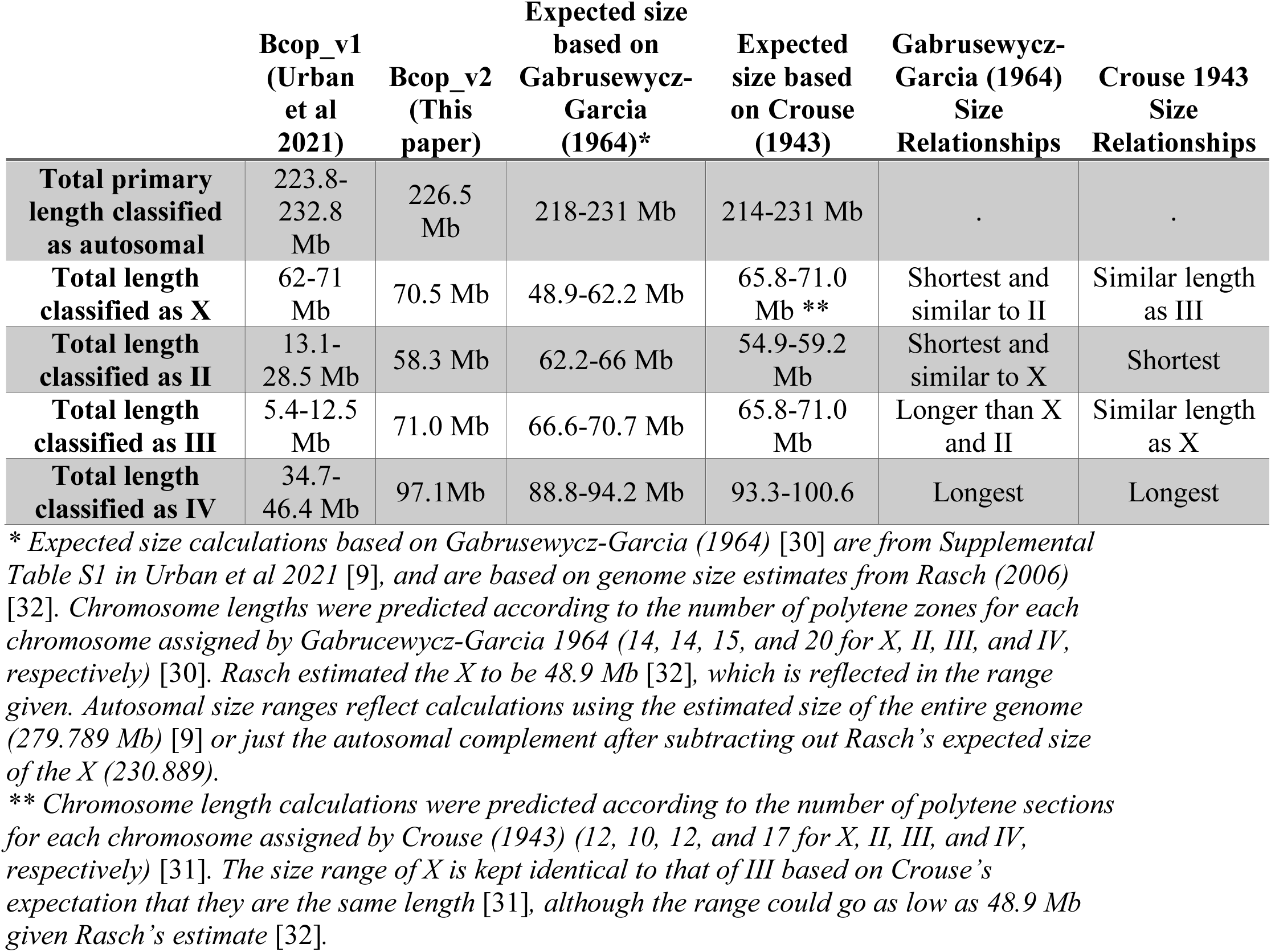
Chromosome anchoring statistics compared to the previous reference assembly.

### Assessing the consistency of the Hi-C data with chromosome specificity and structural accuracy of the scaffolds

With regard to scaffolding accuracy, we interrogated ‘chromosome specificity’, the degree to which contigs in a given scaffold are likely to come from the same chromosome, and ‘structural accuracy’, the degree to which contigs are joined together in the correct order and orientation with respect to the chromosome. Before pursuing orthogonal evaluations of both metrics (see subsequent sections), the Hi-C signal across Bcop_v2 was manually inspected to ensure that these metrics were optimized according to expected interaction frequency patterns.

First, intra-chromosomal interaction frequencies are expected to exceed inter-chromosomal interaction frequencies [11]. As such, for high ‘chromosome specificity’, the same should be true for interaction frequencies within scaffolds compared to between scaffolds. As can be seen in Fig. 2A, the regions of high and low interaction frequencies are all over the map for the input into scaffolding (Bcop_v1_corrected) where contigs are simply ordered from longest to shortest. However, after scaffolding, in the Hi-C map for Bcop_v2, the contigs are ordered such that those regions of high average interaction frequencies are confined within four chromosome-sized scaffold squares along the diagonal and the regions of lowest interaction frequency correspond to inter-scaffold contacts (Fig. 2B). This indicates that the contigs within a given scaffold correspond to the same chromosome and that different scaffolds correspond to different chromosomes.

Second, within a chromosome, interaction frequencies are expected to be highest between adjacent loci and to decay with distance along the linear chromosome sequence [11]. As such, for high ‘structural accuracy’ within a given scaffold, short distances should have the highest interaction frequencies both within contigs and also between adjacent ends of neighboring contigs within the scaffold. The former was ensured in the previous pre-scaffolding “correction” step and the latter will hold true if the contigs were ordered and oriented correctly. High structural accuracy of Bcop_v2 is indicated by the strong Hi-C interaction frequency signal along the diagonal within the chromosome scaffolds (Fig. 2C-F), with no disruptions between neighboring contigs, and by the weaker off-diagonal interaction frequencies that rapidly decay with distance. In addition, several contigs were selected for further analysis of the interaction frequencies connecting them to adjacent contigs. For the start (or end) of each contig inspected, there were orders of magnitude more interactions with the immediately adjacent terminal of the nearest neighboring contig than any other region of any other contig within the scaffold. It also had comparable interactions with its opposite flank within the same contig (see example in Fig. 2G). Overall, these results indicate that the adjacent contigs within a given scaffold correspond to adjacent loci in the chromosome according to the logic of Hi-C.

In sum, the Hi-C signal is consistent with well put together chromosome scaffolds that have both high chromosome specificity of the contigs within each scaffold and high structural accuracy in how they are ordered. Nonetheless, since the scaffolds were put together using this same Hi-C logic and data, we sought other ways to test the chromosome specificity and structural accuracy of the chromosome scaffolds, including looking at the order of multiple known anchor sequences, centromere positions, how repeats are distributed across the scaffolds, known features expected for the X, and comparisons to the chromosomes of other Dipteran genomes, all detailed below.

### Establishing the chromosome identity, polytene orientation, and structural accuracy of the chromosome scaffolds using the locations of multiple anchor sequences

The chromosomal identities of the chromosome-scale scaffolds were obtained using “anchor” sequences for 10 known unique chromosomal locations [17–29] by mapping them to the scaffolds (Fig. 3A-D; Tables 2 and 3; see Methods). All anchor sequences from a given chromosome mapped to the same chromosome-scale scaffold, and there were no conflicts of chromosome identity among the multiple anchor sequences (Fig. 3A-D). These results further confirm that each chromosome-scale scaffold corresponds to one and only one chromosome and that each scaffold corresponds to a different chromosome than the three other scaffolds. Thus, the anchor sequences are concordant with the high ‘chromosome specificity’ of the contigs within scaffolds observed in the Hi-C maps.

**Figure 3:**
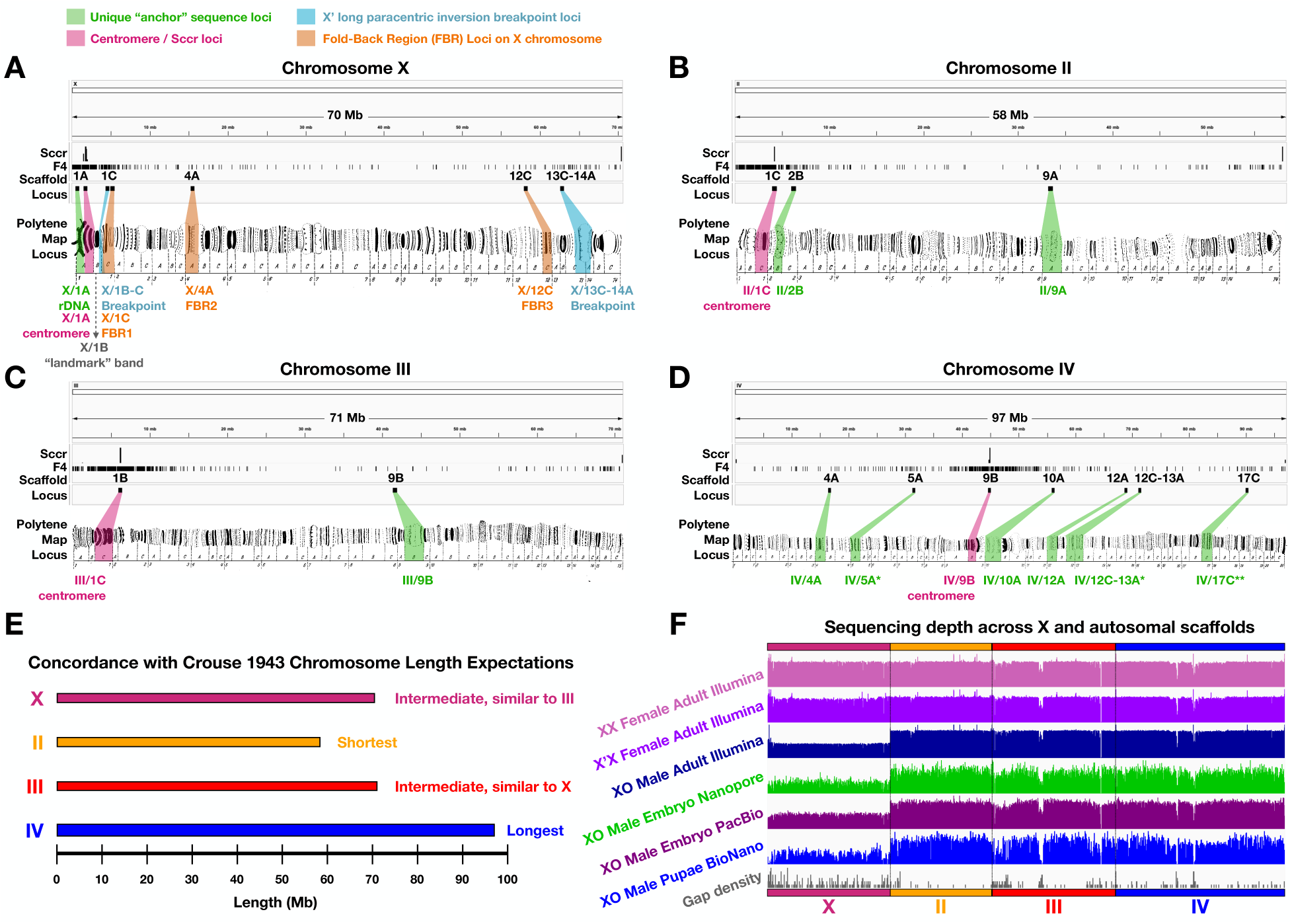
Chromosome identity, orientation, and assessment of the chromosome-scale scaffolds. **(A-D)** Anchoring, polytene map orientation, scaffold structural verification, and loci of interest on the chromosome-scale scaffolds for chromosomes **(A)** X, **(B)** II, **(C)** III, and **(D)** IV. The chromosome identity and specificity of each scaffold, their consistent orientations with polytene maps, and structural verifications were obtained using 10 unique “anchor” sequences with known chromosomal addresses mapped previously using *in situ* hybridization [17–29]. Centromeric sequences identified the expected locations of centromeres [22,30,31,45] on acrocentric (X, II, and III) and meta-centric (IV) chromosomes, which were further supported by the density of a probe sequence (F4 from ScRTE) shown previously to be pericentromeric in all four chromosomes [45]. The scaffold for X also shows the locations found for the fold-back regions and long paracentric inversion breakpoints on the X’. Each chromosome-scale scaffold had at least one known unique sequence location anchoring it to a specific chromosome. There was no conflicting evidence with regard to chromosome identity nor with regard to the order in which sub-chromosomal sequences mapped along the scaffolds. Polytene chromosome maps were reproduced from plates 1, 2 and 3 of Gabrusewycz-Garcia (1964) [30] with permission from Springer Nature under permission number: 5490830617309. **(E)** Chromosome scaffold sizes are consistent with chromosome length expectations from Crouse (1943) [31]. **(F)** Chromosome scaffolds are consistent with read depth expectations on the X versus autosomes in males and females. Overall, this evidence suggests that Hi-C accurately ordered and oriented the contigs into chromosome-scale scaffolds corresponding to the expected four somatic chromosomes.

**Table 3:**
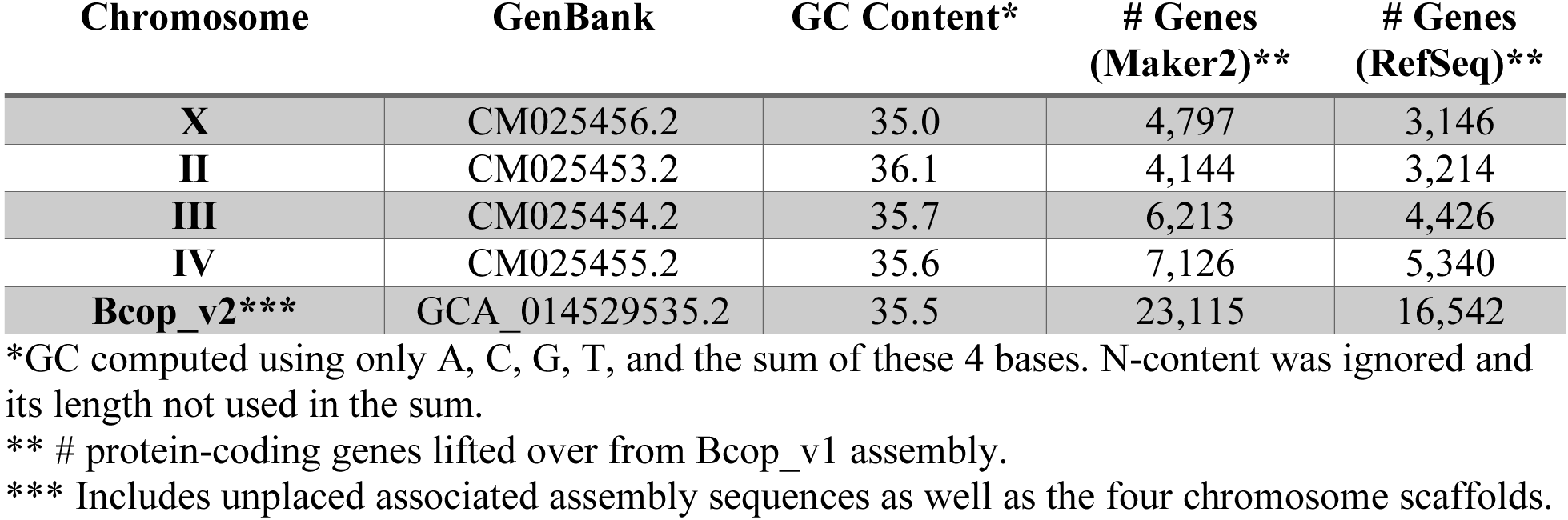
Chromosome-scale scaffold accessions and attributes.

The structural accuracy was further tested by the analyzing the order of the anchor sequences across the scaffolds in addition to the approximate positions of centromeres mapped onto each scaffold with a known centromeric sequence (Fig. 3A-D; see below Results and Methods). Since both the polytene map and the scaffold locations of the 10 anchor sequences and 4 centromeres were known, it was also possible to orient the chromosome scaffold sequences in the same direction as locus numbers have been assigned on polytene maps for all four chromosomes [30] (Fig. 3A-D), which was the final processing step for Bcop_v2. Importantly in terms of the structural integrity of how the scaffolds were ordered by Hi-C, when multiple anchors were present for a given chromosome, they all mapped in the expected order along the same scaffold (Fig. 3A-D). Thus, a sampling of 14 sites across the four chromosomes further confirms the high structural accuracy of the order of contigs along the chromosome-scale scaffolds.

### Chromosome lengths and copy numbers further confirm chromosome identities

The chromosome identities of each scaffold were further supported by comparing the scaffold lengths with prior expectations of the chromosome lengths. In very early studies of these chromosomes by H.V. Crouse [31], chromosome II was predicted to be shortest, chromosome IV was established as the longest, and chromosomes X and III were expected to be of equivalent lengths. Crouse’s early expectations are confirmed by the chromosome-scale scaffolds (Table 2; Fig. 3E), of which II is shortest (58.3 Mb), IV is the longest (97.1 Mb), and X and III are approximately the same length (70.5 and 71.0 Mb, respectively). Moreover, the chromosome lengths reflected by the scaffolds are concordant with the expected sizes of each chromosome using the number of polytene sections assigned by Crouse [31] and the expected genome size from DNA content measurements [32] (Table 2). The total length (226 Mb) of the autosomal scaffolds (II, III, IV) was also concordant with these predictions (214-231 Mb) (Table 2). Note that the chromosome length expectations and the number of zones in each from Crouse [31] were later revised such that X and II were both shortest and of equivalent length [30], but this under-estimate of the X was anticipated by Crouse [31] and is now refuted by the chromosome-scale scaffolds that support Crouse’s earlier length expectations (Table 2; Fig. 3E).

The chromosome identity of the X chromosome scaffold and of the autosomal nature of the other scaffolds (II, III, and IV) were further supported by read depth analysis. Females are diploid for both the autosomes and the X chromosome. Males are diploid for the autosomes, but haploid for X. Thus, read depth for females along the X and autosomes should be equivalent whereas read depth along the X from males should be half that of autosomes. These read depth expectations are borne out by multiple genomic technologies (Fig. 3F), including paired-end Illumina reads for both types of adult female and adult males [10], long reads from Nanopore and PacBio for male embryos [9]; and BioNano Genomic optical maps from male pupae [9]. The diploid-level read depth in both males and females across the scaffolds for chromosomes II, III, and IV confirms that they are strictly composed of autosomal contig sequences (Fig. 3F). Similarly, the haploid-level read depth in all male samples, but diploid level in female samples, across the entirety of the X chromosome scaffold, confirms not only the chromosome identity of the scaffold as X, but that all the contigs within it are indeed X-linked (Fig. 3F).

### Gene annotations for the chromosome-scale assembly (Bcop_v2)

There are presently two high quality gene annotation sets produced using the previous genome assembly (Bcop_v1). One was constructed by us [9] using Maker2 [33], which can be found at USDA Ag Data Commons [34]. The other was made by the NCBI Eukaryotic Genome Annotation Pipeline for RefSeq, is called NCBI *Bradysia coprophila* Annotation Release 100, and can be found at NCBI [35]. These gene annotations have already been demonstrated to score highly in ‘completeness’ as determined by the number of Dipteran BUSCOs (97%) found within them, where the BUSCOs used were defined as genes present as single copy orthologs (SCOs) in 95% of Dipterans [9,36]. While the FASTA files containing the transcript and protein sequences are in no need of updating, the GFF files that detail the coordinates of the gene models, including their exons, introns, CDS, and UTRs, on Bcop_v1 are not useful for the updated assembly, Bcop_v2. Since Bcop_v2 was produced by stitching together the Bcop_v1 sequences, a simple “liftover” procedure, where features from an assembly can be exactly mapped to an updated version, was adequate for creating new GFF files detailing the coordinates of the genes on Bcop_v2.

Both gene annotation sets had similar lift-over success rates for all metrics (Table 4). Using the Maker2 annotation set as an example, of the 23,117 genes and 28,870 transcripts, only 2 genes with 4 transcripts were not mapped to Bcop_v2. Of the 28,866 transcript models successfully lifted over to Bcop_v2, all but one (28,865) had perfect full-length alignments to the original transcript sequences, although 52 (<0.2%) were a bit shorter than the original. In total, only 53 transcript sequences had differences in the updated genome assembly (Table 4). Even fewer, just 49 transcripts, had differences at the level of protein sequence, which is to say that nearly all (99.83%) had identical protein sequences between assemblies (Table 4). An even higher percent at the gene level was identical between assemblies. For example, 99.87% of lifted-over genes had identical protein sequences for all transcript isoforms, and 99.91% had identical protein sequences for all or at least a subset of transcript isoforms (Table 4).

**Table 4:** Lifted-over gene annotation statistics.

|  | **Maker2**<br>**(Urban et al**<br>**2021)\*** | **NCBI r100**<br>**(RefSeq)\*\*** |
| --- | --- | --- |
| **# protein-coding genes Bcop_v1** | 23,117 | 16,546 |
| **# genes lifted over to Bcop_v2** | 23,115<br>(99.99%) | 16,542<br>(99.98%) |
| **# genes not lifted over to Bcop_v2** | 2<br>(0.01%) | 4<br>(0.02%) |
| **# transcripts Bcop_v1** | 28,870 | 28,473 |
| **# transcripts lifted over to Bcop_v2** | 28,866<br>(99.99%) | 28,464<br>(99.97%) |
| **# transcripts not lifted over to Bcop_v2** | 4 (0.01%) | 9 (0.03%) |
| **# lifted over transcript sequences identical after liftover** | 28,813<br>(99.82%) | 28,388<br>(99.73%) |
| **# lifted over transcripts that yield identical protein sequence**<br>**after liftover** | 28,817<br>(99.83%) | 28,396<br>(99.76%) |
| **# lifted over transcript sequences changed after liftover** | 53<br>(0.18%) | 76<br>(0.27%) |
| **# lifted over transcripts that yield changed protein sequence**<br>**after liftover** | 49<br>(0.17%) | 68<br>(0.24%) |
| **# lifted over genes where all transcript isoform sequences are**<br>**identical before and after liftover** | 23,083<br>(99.86%) | 16,512<br>(99.82%) |
| **# lifted over genes where all transcript isoforms yield same**<br>**protein sequences after liftover** | 23084<br>(99.87%) | 16516<br>(99.84%) |
| **# lifted over genes where a subset of transcript isoforms**<br>**sequences is identical before and after liftover** | 10<br>(0.04%) | 3<br>(0.02%) |
| **# lifted over genes where a subset of transcript isoforms yield**<br>**same protein sequences after liftover** | 10<br>(0.04%) | 4<br>(0.02%) |
| **# lifted over genes where all transcript isoform sequences**<br>**differ before and after liftover** | 22<br>(0.1%) | 27<br>(0.16%) |
| **# lifted over genes where all transcript isoforms yield changed**<br>**protein sequences after liftover** | 21<br>(0.09%) | 22<br>(0.13%) |

\* Urban et al (2021) [9,34]
\*\* NCBI *Bradysia coprophila* Annotation Release 100 [9,35]

We also visually confirmed in a genome browser that gene models at the same loci look the same between assemblies. Moreover, the lifted over gene models were highly concordant with RNA-seq coverage as illustrated in Fig. 4A-F, which shows six example genes of interest from recent *B. coprophila* papers [7,37] and that, for simplicity, shows read depth from only one RNA-seq sample of irradiated larvae [37]. The gene models were highly consistent with RNA-seq coverage from all datasets we checked, including from various life stages [9], radiation conditions [37], and unpublished data. Overall, gene/protein sequence comparisons between assemblies and gene model concordance with RNA-seq both demonstrate that the lift-over process successfully transferred the annotation information from Bcop_v1 to Bcop_v2. The chromosomal location of genes allowed us to analyze the number of genes per chromosome (Table 3) as well as chromosome-specific log2 fold-changes of male versus female gene expression in multiple life stages (Fig. 4G), which supported a dosage compensation mechanism on the single male X as we and others found previously [9,38].

**Figure 4:**
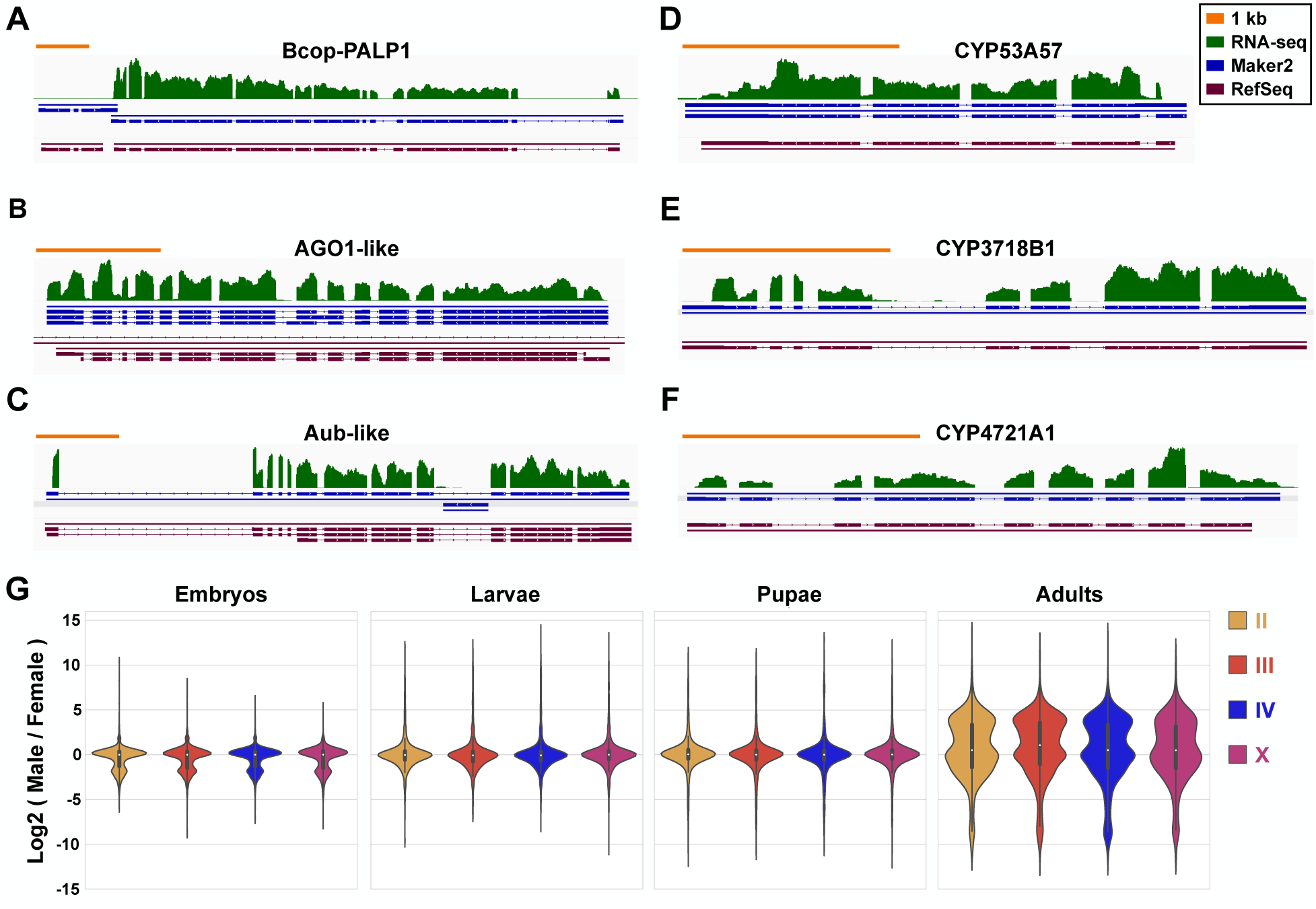
Analysis of gene models on Bcop_v2 lifted over from Bcop_v1 gene annotation sets. **(A-F)** Example Bcop_v1 gene models were chosen to show concordance with RNA-seq datasets after liftover to Bcop_v2. For simplicity, only one RNA-seq dataset is shown (irradiated fourth instar pre-eyespot larvae, replicate 1 [37]). However, lifted over gene models were consistent with all RNA-seq datasets checked, including those from all life stages and both sexes used in the original gene annotation [9]. RNA-seq concordance is illustrated here using genes of interest from recent *B. coprophila* studies, including **(A-C)** PARP and Argonaute family genes that arose in response to radiation [37], and **(D-F)** example “alien” cytochrome P450 genes [7]. For all six gene panels **(A-F)**, as summarized in the pictorial legend box on the top right: the orange scale bar represents 1 kb, the green trace (top level) shows RNA-seq coverage, the blue gene models (middle level) are from Maker2 [9,34], and the purple gene models (bottom level) are from NCBI RefSeq [9,35]. In all cases, the RNA-seq trace shows genome-wide coverage that was computed agnostic to the location of gene structures, and that was not artificially restricted to exons. Thus, the correspondence of higher RNA-seq coverage over exons shows that fidelity of the liftover process. **(G)** NCBI gene models lifted over to Bcop_v2 were used for chromosome-specific differential expression analysis of RNA-seq data from males versus females across four life stages [9]. Log2 fold-changes are shown for genes grouped by chromosome-scale scaffolds. The distribution of male vs. female fold-changes across X-linked genes match the distributions across each autosomal chromosome rather having a 2-fold down-shifted X-linked fold-change distribution corresponding to the single male X chromosome compared to two in females. This indicates there is dosage compensation.

### Gene sets on chromosome scaffolds fit expectations of chromosome similarity with other species as a function of evolutionary distance

Over evolutionary time, chromosomes tend to undergo rearrangements that include translocations. Through these rearrangements, genes get shuffled around the genome, and more gene shuffling is likely to occur with increasing evolutionary distance. When defining chromosomes as sets of genes, two closely related species are likely to have very similar chromosomes and therefore highly intersecting chromosomal gene sets. In contrast, distantly related species are likely to have smaller intersections between chromosomal gene sets, approaching “random” proportions of shared genes between arbitrarily selected chromosomes from each species. While seeing this trend with respect to *B. coprophila* genes on Bcop_v2 chromosome-scale scaffolds would further support the accuracy of the Hi-C scaffolding process, not seeing this trend would raise concerns.

To test the evolutionary prediction described above, the chromosomal groupings of *B. coprophila* genes (chromosomal gene sets) was compared with that of four other fly (order Diptera) species with known evolutionary relationships (Fig. 5A) and which all had chromosome-scale genome assemblies [39–42]. The most distantly related species selected was *Drosophila melanogaster* from the so-called “higher Diptera” (sub-order Brachycera). In contrast, *B. coprophila* and the three remaining species are all part of the “lower Diptera” (sub-order Nematocera), which diverged from the higher Diptera ∼200 MYA [43]. The closest related species to *B. coprophila* was *Pseudolycoriella hygida*, which is within the same family (Sciaridae) commonly known as black fungus gnats or dark-winged fungus gnats. *P. hygida* had a chromosome-scale genome release in 2023 [42] following the release of Bcop_v2, which was opportune for this analysis. The remaining two Nematocerans are mosquitoes (Culicidae family): the yellow fever mosquito (Aedes aegypti) and the African malaria mosquito (Anopheles gambiae). The proteomes of all fives species were compared to determine orthologs as well as to construct a proteome-wide rooted species tree, which recapitulated the known evolutionary relationships (Fig. 5B). This indicates that the genes determined to be orthologs across species should also be informative when comparing their chromosomal locations.

**Figure 5.**
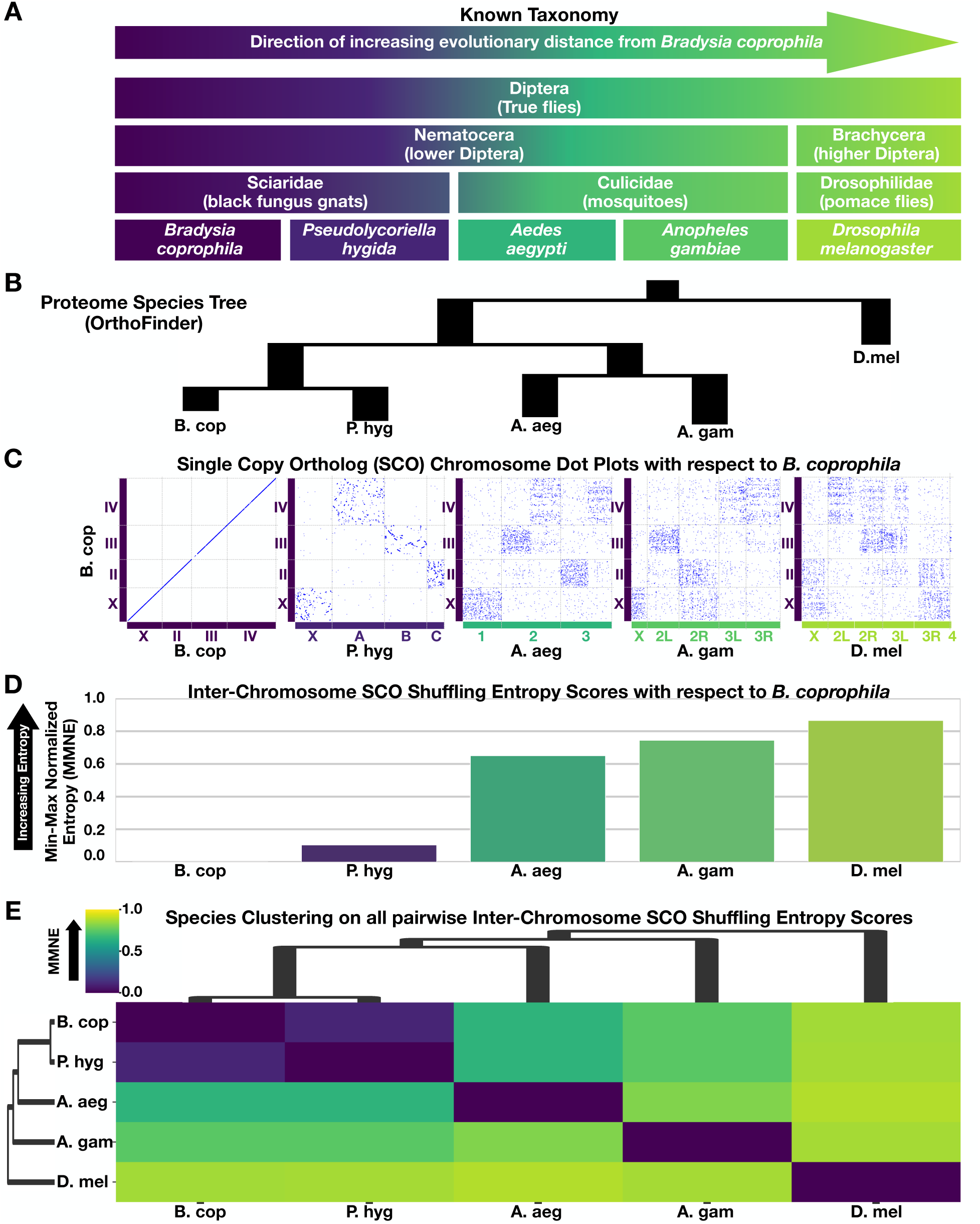
Interchromosomal gene shuffling increases with evolutionary distance, further supporting integrity of *B. coprophila* chromosome-scale scaffolds. **(A)** Known taxonomic and phylogenetic information regarding *B. coprophila* and four other Dipteran species. With respect to *B. coprophila,* the species are presented here from left to right in order of the most closely related to most distantly related. The color scheme from left to right (dark purple to light yellow-green) reflects the entropy scores between *B. coprophila* and these species as in C-E. The species are presented in this order in each sub-figure below as well. **(B)** Species tree produced by OrthoFinder [81,82] using the proteomes of these five species reproduces the expected relationships amongst these Dipteran flies. **(C)** Scatter plots of SCO locations along the chromosomes of each of the five species (x-axis) plotted against *B. coprophila* (y-axis). Inter-chromosomal SCO scattering increases with evolutionary distance as expected. Blue dots represent SCO locations. Colored boxes along x- and y-axes represent chromosomes. The chromosome box color scheme reflects the entropy score color scheme in D-E. **(D)** The pairwise min-max normalized entropy scores for *B. coprophila* computed against all five species. The entropy scores increase with evolutionary distance as expected. Although the entropy scores approach a fully random state with evolutionary distance from *B. coprophila*, pairwise relationships of chromosomal SCO groupings still display a bit of non-randomness even in the most distantly related fly species tested (*D. melanogaster*). **(E)** Matrix of all pairwise min-max normalized entropy (MMNE) scores among the five species presented as a hierarchically-clustered heatmap. The dendrogram from pairwise entropy scores reproduces the expected relationships among these Dipteran flies. Altogether, as the SCO dispersion and entropy of chromosomal gene sets follows evolutionary expectations, the location of genes in the chromosome-scale scaffolds we report here for *B. coprophila* are non-random and suggest the contigs were grouped correctly by chromosome.

To test if the degree of gene shuffling around the chromosomes of these five species also reflected their known phylogenetic relationships, the genomic locations of “Single copy orthologs” (SCOs) were visually compared across all five species (Fig. 5C) with respect to their locations within *B. coprophila* chromosomes (Bcop_v2). The five species had 4,297 SCOs. The chromosomal addresses of SCOs were compared between each pair of species relative to *B. coprophila* in order to (i) map *B. coprophila* chromosomes to their most likely counterparts in each other species (Table 5) and (ii) visualize as dot plots (Fig. 5C). *B. coprophila* was plotted against itself to demonstrate the base case of what the absence of gene shuffling looks like (Fig. 5C). Visually, the amount of gene shuffling increased with evolutionary distance as expected. While there is gene shuffling even between *B. coprophila* and the closest related species tested (*P. hygida*), it is mostly intra-chromosomal and this pair demonstrates the highest amount of conservation of SCOs from given *B. coprophila* chromosomes to their single corresponding chromosomes in the other species (Fig. 5C; Table 5). In contrast, the most distantly related species (*D. melanogaster*) visually has the most gene shuffling across chromosomes, and the mosquito species have intermediate levels, as expected (Fig. 5C; Table 5). Thus, visual inspection of the pairwise SCO dot plots makes a strong case that the *B. coprophila* gene sets defined by the chromosome-scale scaffolds (Bcop_v2) follow evolutionary expectations.

**Table 5:** Chromosomal counterparts in other species.

**Chromosome(s) from other species with highest percent of SCO from given *B. coprophila* chromosome**
| *B. coprophila*<br>chromosomes | *P. hygida* | *A. aegypti* | *A. gambiae* | *D. melanogaster* |
| --- | --- | --- | --- | --- |
| **X** | X (97.3%) | 1 (83.5%) | 2 (54.3%);<br>X (34.8%) | 3 (57.1%);<br>X (29.7%) |
| **II** | C (97.3%) | 3 (81.9%) | 2 (84.2%) | 3 (59.0%);<br>X (27.9%) |
| **III** | B (97.0%) | 2 (85.1%) | 2 (80.0%) | 3 (47.2%);<br>2 (45.2%) |
| **IV** | A (98.2%) | 2 (50.7%);<br>3 (44.8%) | 3 (81.8%) | 2 (70.3%);<br>3 (23.9%) |

The amount of “gene shuffling” was also quantified between all pairs of the five species using the single summary metric of “entropy”. Briefly, entropy can be thought of as a measure of “mixed-up-ness” that can quantify systems as they evolve from a non-random to a random state [44]. Here, it is used to measure the amount of inter-chromosomal gene shuffling between pairs of species, and ultimately as a function of evolutionary distance from *B. coprophila*. A normalized entropy metric that ranges between 0 and 1, called Min-Max Normalized Entropy (MMNE), was computed for each pairwise comparison of two species. MMNE shows the observed entropy relative to the minimum and maximum entropy values possible for that pairwise species comparison given the number of shared SCOs, chromosomes, and associated probabilities (see Methods; see example SCO probabilities in Table 6). The initial non-random lowest entropy state (when MMNE is 0) is simply a comparison to self (Fig. 5D). Entropy is expected to increase (MMNE approaches 1) with more distantly related species that had larger periods of time for inter-chromosomal rearrangements to shuffle genes around, which is what is observed with increasing evolutionary distance from *B. coprophila* (Fig. 5D). Furthermore, performing clustering on the matrix of all pairwise inter-species entropy scores reproduces the species groupings expected by the known evolutionary relationships (Fig. 5E), putting the fungus gnat family members (*B. coprophila* and *P. hygida*) together within the larger Nematoceran cluster that also included the pair of mosquitoes, all of which were on a separate branch from the single Brachyceran*, D. melanogaster*. The results of these comparative genomic evaluations further bolster the confidence we have in the Hi-C directed process that yielded Bcop_v2 chromosome-scale scaffolds.

**Table 6:** Joint probabilities of SCOs on chromosomes from *B. coprophila* and *P. hygida* as an example. This is an example, showcasing the two fungus gnats, of the input used for computing the raw entropy score before min-max normalization (see Methods), including the observed marginal probabilities of finding a single copy ortholog (SCO) on a given chromosome from a given species and the observed joint probabilities of finding a SCO on a given pair of chromosomes between the species. *B. coprophila* and *P. hygida* chromosomes are ordered by corresponding chromosomes as defined in Table 5, such that the highest joint probabilities are along the diagonal (left-to-right, top-to-bottom). The joint probabilities correspond to the proportions of SCOs that appear on (or the probabilities that any given SCO will appear on) each pair of inter-species chromosomes. Marginal probabilities (italicized) are obtained by summing the non-rounded values in the rows for *B. coprophila* and the columns for *P. hygida* (summed values rounded to 3 digits)*. The raw entropy score is computed on the observed joint probabilities. To get the minimum possible entropy the joint probabilities between these species could take given their SCO distributions, entropy is computed on each set of marginal probabilities separately, then averaged. To get the maximum entropy, entropy is computed on the set of joint probabilities expected at random, which are defined as the products of each inter-species pair of marginal probabilities. Min-max normalized entropy (MMNE) scores are then computed as: (observed entropy – minimum entropy)/(maximum entropy – minimum entropy). See the Methods for further details. * *Note that joint probabilities were rounded to 3 digits for simplifying this table, but not for computing the entropy. This results in a difference between the sum of rounded joint probabilities in a column or row and the reported marginal probability computed from summing non-rounded joint probabilities, then rounding the sum. The difference is only in the third position after the decimal*.

|  |  | *P. hygida* chromosomes |  |  |  |  |
| --- | --- | --- | --- | --- | --- | --- |
|  |  | X | C | B | A | *Marginal*<br>*B. cop* |
| *B. coprophila*<br>chromosomes | X | 0.253 | 0.002 | 0.002 | 0.003 | *0.261* |
|  | II | 0.000 | 0.166 | 0.001 | 0.003 | *0.170* |
|  | III | 0.002 | 0.001 | 0.241 | 0.004 | *0.248* |
|  | IV | 0.002 | 0.002 | 0.002 | 0.315 | *0.321* |
| *Marginal*<br>*P. hyg* |  | *0.258* | *0.170* | *0.247* | *0.325* |  |

### Investigating reported horizontal gene transfer using the chromosome scaffolds, Hi-C, and other genomic data

Recently an analysis of the CYPome (P450 genes) of *B. coprophila* lead to the conclusion that there has been horizontal gene transfer (HGT) from springtails, mites, and fungi [7]. There were several lines of evidence that these so-called “alien” genes, which appear to have been obtained by HGT, are truly integrated into the nuclear chromosomes as opposed to being present from contamination sources [7]. For example, the “alien” genes had homology and synteny in the genomes of closely related species from three different continents and had evidence of being expressed and spliced [7], which is also shown for three such genes in Fig. 4. The locations of these genes in the chromosome-scale scaffolds allowed a further investigation into the likeliness that these ‘alien’ genes are truly integrated into the chromosomes of *B. coprophila* with a variety of orthogonal genomic datasets, including Hi-C, long reads, and optical maps. First, the locations of the P450 genes in the chromosome-scale scaffolds did not give rise to disrupted or otherwise unexpected Hi-C signals. The contigs they reside on survived the Hi-C guided correction step and were stitched into the chromosome-scale scaffolds as part of the Hi-C guided scaffolding step. Moreover, the alien P450 genes have typical Hi-C interaction frequencies that decay with distance that are comparable in all cases to nearby ‘highly conserved fly genes’ of similar length (Fig. 6). Second, across a variety of genomic datasets from different technologies and samples, there were no disruptions in sequencing depth seen over regions containing the putative “alien” P450 genes (Fig. 6) whereas disruptions were seen over gaps in the assembly as expected. Finally, there exist Hi-C interactions and single DNA molecules corresponding to long reads and optical maps that physically connect the alien P450 genes to highly conserved Dipteran genes (data not shown). In summary, while we make no claims here about HGT, we do confirm that the so-called “alien” P450 gene sequences in the *B. coprophila* genome assembly are further supported as being part of the nuclear chromosomes rather than from a separate contamination source by our analyses of a diverse set of genomic datasets. The same conclusion can be made from similar analyses of five terpene synthase genes within the *B. coprophila* chromosome-scale scaffolds (Bcop_v2), which also have been proposed to have originated by HGT from mites [8].

**Figure 6:**
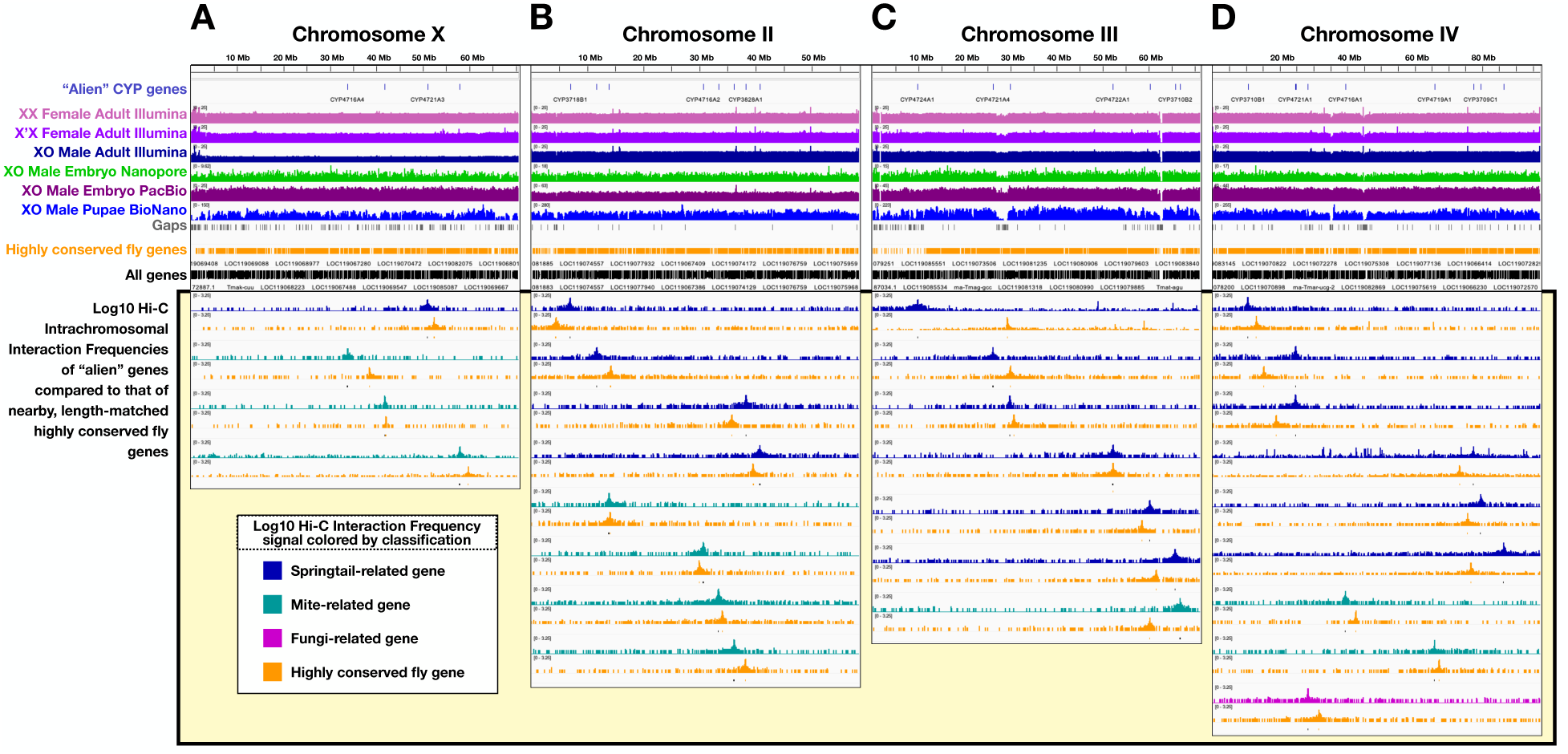
Diverse genomic datasets support the existence of “alien” genes within the chromosomes. IGV traces for chromosome **(A)** X, **(B)** II, **(C)** III, and **(D)** IV. From the top track downward, each shows the location of (1) “alien” CYP genes described by Feyersen et al [7], (2-7) coverage tracks from various genomic technologies and samples [9,10], (8) gaps in the assembly where coverage signal is expected to drop, (9) locations of all highly conserved fly genes, and (10) locations of all genes. In each, the 11^th^ track from the top and all tracks beneath it highlighted by the yellow box correspond to the log10 intrachromosomal interaction frequency (IIF) signal emanating from selected “alien” genes (dark blue, cyan, or magenta) or the closest by highly conserved fly gene that is approximately the same length (orange). All log10 IIF tracks have the same y-axis range (0-3.25), allowing direct comparison of peak heights and shapes. Under each pair of log10 IIF signals is a track showing the exact location of both genes. Overall, the genes reported to be “aliens” by Feyersen et al [7] show no differences in coverage or Hi-C interactions compared to highly conserved fly genes, demonstrating their existence inside the scaffolds is not a result of contamination and/or assembly errors.

### The locations of known centromeric repeats and peri-centromeric transposons further support the chromosomal structures of the scaffolds

Centromere locations have previously been mapped on polytene maps [22,30,31,45]. Centromere positions in all chromosome scaffolds were approximated using the centromere-associated repeat sequence, Sccr (*<u>S</u>ciara <u>c</u>oprophila* <u>c</u>entromeric <u>r</u>epeat), that is known to hybridize to the centromeres of all four somatic chromosomes [45], and an associated repeat family that contains Sccr (see Methods). The Sccr-repeat-family-enriched positions were found at the expected centromeric locations in all chromosome scaffolds (Fig. 3A-D, Fig. 7A). Specifically, chromosomes X, II, and III are acrocentric with centromeres very close to the beginning of these chromosomes (as oriented) at polytene zones X/1A, II/1C, and III/1C. In agreement, Sccr mapped within the first few megabases of the corresponding 58-71 Mb scaffolds (Fig. 3A-D, Fig. 7A). In contrast, chromosome IV is metacentric with a centrally-located centromere at polytene zone IV/9B, where zones span from IV/1A to IV/20C. As expected, Sccr mapped to a medial region, at positions spanning ∼44.7-45.0 Mb, of the 97.1 Mb chromosome IV scaffold (Fig. 3A-D, Fig. 7A).

**Figure 7:**
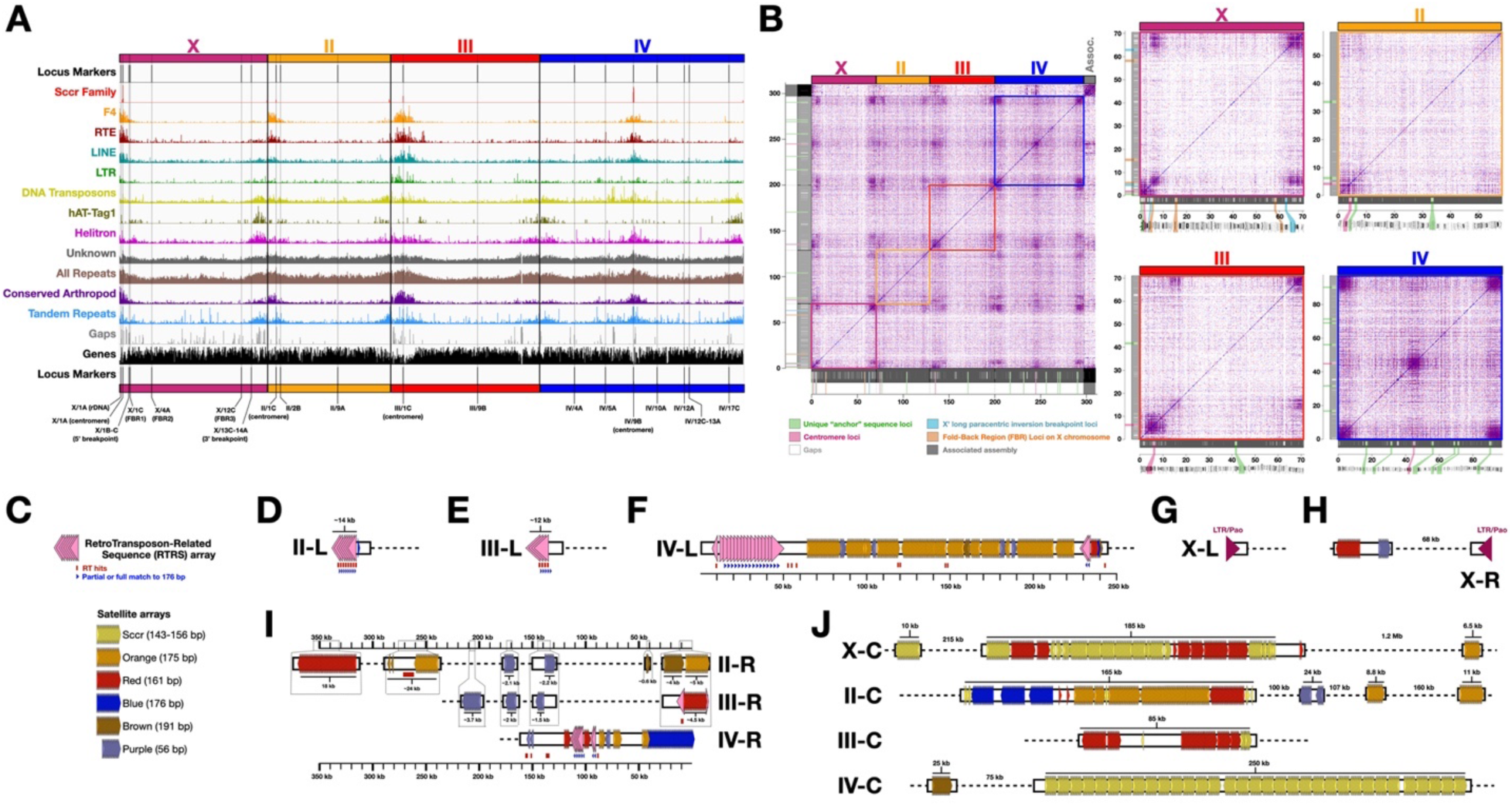
Repeat regions, centromere locations, and terminal repeats across the *B. coprophila* chromosome-scale scaffolds. **(A)** Repeat densities in 100 kb bins across each chromosome, and locations of known locus markers. Sccr = Sciara coprophila centromeric repeat [45]. F4 = probe sequence for ScRTE [45]. hAT-Tag1 = a DNA transposon sub-family. RTE = a LINE sub-family. F4, RTE, LINE, and LTR are retrotransposon families. Helitron elements are DNA transposons thought to amplify through a rolling-circle mechanism. Unknown = repeats that were modeled but not classified as a known repeat class. All Repeats = all in comprehensive repeat library. Conserved Arthropod = matches to known Arthropod Repeats (excluding *B. coprophila*). Tandem Repeats were found with Tandem Repeats Finder [89]. Gaps = assembly gaps. **(B)** Dot plots corresponding to (i) the entirety of Bcop_v2, including all primary chromosome scaffolds as well as the associated contigs, and (ii) each individual chromosome mapped against itself. The locations of various loci markers are shown in the margins, are color-coded as shown in the legend, and are extended to axes. Gap locations are shown as white rectangles in the grey margins. Polytene chromosome maps were reproduced from plates 1, 2 and 3 of Gabrusewycz-Garcia (1964) [30] with permission from Springer Nature under permission number: 5490830617309. **(C)** Legend to help interpret D-J. **(D-G)** Pictorial models of the left termini for scaffolds corresponding to **(D)** chromosome II (II-L), **(E)** chromosome III (III-L), **(F)** chromosome IV (IV-L), and **(G)** chromosome X (X-L). **(H-I)** Pictorial models of the right termini for scaffolds corresponding to **(H)** chromosome X, and **(I)** chromosomes II, III, and IV. **(J)** Pictorial models of satellite sequences in centromeric and pericentromeric regions of each chromosome scaffold. Models represent the approximate positions and lengths of RTRS and satellite arrays. RTRS copy number is shown. Satellite copy number can be approximated by the total length divided by the satellite unit length.

Importantly, the predicted centromeric positions in the chromosome scaffolds were concordant with respect to their relative positions between other anchor sequences (see above Results) in the order of anchor sequence loci along the polytene maps (Fig. 3A-D, Fig. 7A), again highlighting the structural accuracy of the scaffolds.

A specific RTE-like retro-transposable element called ScRTE is increasingly enriched with proximity to all centromeres, which was previously shown experimentally using a sub-sequence from ScRTE, called “F4”, as a probe [45]. If the scaffolds represent the chromosomes well and if the centromeric positions within them defined above using Sccr are correct, then ScRTE should also show increasing enrichment with proximity to those centromeric positions. To test this, we found all matches specifically to the F4 probe sequence across the chromosome scaffolds as well as the distribution of all RTE elements identified during repeat annotation (described below). Both showed the expected trend of RTE transposons being increasingly enriched with proximity to the centromere positions approximated by Sccr (Fig. 3A-D, Fig. 7A). This further supports the centromere positions within the scaffolds. Moreover, it indicates that the contigs were ordered within the scaffolds such that they are increasingly enriched for RTE with proximity to the centromeres, and therefore again signifies high structural accuracy.

### Repeat locations show family-specific biases towards peri-centromeric and sub-telomeric regions

In general, repeats and transposons are often enriched near centromeric, peri-centromeric, telomeric- and sub-telomeric regions in Dipteran chromosomes [39,40,46]. To test this expectation, we analyzed the distribution of repeats across the *B. coprophila* chromosome-scale scaffolds. Centromeric positions are defined here by the positions mapped by the Sccr-associated repeat family as above and peri-centromeric positions are defined as the much broader repeat-rich regions flanking each side. The telomeric and sub-telomeric regions are defined here simply as the regions near and including the scaffold ends, which have been shown to correspond to sub-telomeres and telomeres in other Hi-C-based scaffolds of insect assemblies [47,48]. If present, telomeric repeats are expected only in the terminal-most 10-50 kb, and sub-telomeres are the immediately-adjacent, much broader repeat-rich regions at chromosome ends. For the sake of clarity, and to avoid conflating telomeres and sub-telomeres, the beginning and end of each chromosome scaffold, as oriented, will sometimes be referred to as its left and right termini, respectively.

First, the entire genome assembly was mapped against itself (or individual chromosome scaffolds against themselves) to create dotplots that naively show all pairwise regions that share high sequence similarity with each other (Fig. 7B). The regions around and including centromeric positions and scaffold termini were very obvious in these plots, with highly repetitive natures locally within each chromosome scaffold and also with high similarity among the corresponding regions across all chromosome scaffolds. The first 10 Mb of the left terminus of chromosome X is composed of two adjacent repeat blocks. The first repeat block at the beginning of X corresponds to the left telomeric/sub-telomeric zone, rDNA, and centromeric repeats (Sccr) among other centromere-proximal repeats (Fig. 7B). To the right of the centromere but still on the left side of the chromosome, the second repeat block contains two adjacent loci of interest that are discussed in a subsequent section (centromere-proximal X’ breakpoint and putative “FBR1”), and is more similar to the repeat block in the final 5-8 Mb of the right terminus of the scaffold (distal telomeric side) than it is to the left terminal block (Fig. 7B). The chromosome III scaffold has a similar structure on its left terminus. It starts with a 10-12 Mb, centromere-containing, high density repeat block followed on its right by a second smaller repeat block that is more closely related to the terminal ∼5 Mb repeat block at the distal telomeric end on the right terminus of the scaffold than the centromeric end to the left terminus (Fig. 7B). Unlike X and III, the scaffold for chromosome II has a single large (∼8 Mb) repeat block at its centromeric end (left terminus) that has high similarity with the terminal ∼8 Mb at its distal telomeric end on its right side (Fig. 7B). The repeat densities of the chromosome II repeat blocks seem lighter than the other chromosomes in this method of analysis, but not necessarily so in other analyses (Fig. 7A-B). The metacentric chromosome IV scaffold has a repeat block at both termini, each ∼8 Mb, as well as a repeat block in the middle of ∼10 Mb surrounding the approximated centromere position in the scaffold (Fig. 7B). Finally, the sequences in the “associated assembly”, which are not placed in the chromosome-scale scaffolds, are enriched for matches within the centromere-proximal repeat blocks, but also contain matches to rDNA and rDNA-proximal repeats, and matches to terminal repeat blocks to a lesser extent (Fig. 7B). Overall, the repeat structures of the scaffolds help confirm their centromere regions and follow expectations of repetitive structures associated with scaffold termini corresponding to telomeric and/or sub-telomeric regions.

To further analyze the distribution of repeats in the chromosome-scale scaffolds, major repeat families were modeled, annotated, classified when possible, and their relative densities visualized along the linear scaffold sequences (Table 7). Tandem repeats were also mapped independently. Both tandem repeats as well as the summation of all modeled repeat families are indeed enriched around defined centromere positions and scaffold termini (Table 7; Fig. 7A; see Methods). When inspecting different transposon families, their densities near centromeres and scaffold termini (a proxy for telomeres) had different biases. Retrotransposable element families (RTE, LTR, LINE) were generally more enriched around centromeric positions than near scaffold termini (Fig. 7A). Their densities are higher at the centromeric (left) ends of the acrocentric chromosomes (X, II, and III) compared to the right ends, and higher at the medial centromere position of IV compared to its termini (Fig. 7A). These families have relatively little or no enrichment in terminal regions, which have similar densities compared to the rest of the chromosome lengths (Fig. 7A). LTR elements are particularly enriched near the centromere of chromosome III (Fig. 7A). Sequence labeled as LINE was most abundant in general (Table 7; Fig. 7A). DNA transposon families have a different bias, seeming to have higher relative densities at least in some telomeric and/or sub-telomeric regions than near centromeres (Fig. 7A). For example, see the non-centromeric ends (right termini) of the scaffolds for chromosomes X and II (Fig. 7A). A sub-family called hAT-Tag1 is a great example of an element whose highest densities are in terminal regions compared to other regions of the chromosome scaffolds, including centromeres (Fig. 7A). Nevertheless, although DNA elements may be more biased in termini of X and II than their centromeres, DNA transposons are also found near all centromeric and peri-centromeric regions, illustrated well by the medial centromere of chromosome IV (Fig. 7A). Moreover, the bias of DNA transposons to these repeat-rich regions compared to gene-rich intervening regions is not as high as for other transposon families (Fig. 7A). Finally, the “Rolling Circle” Helitron family is distributed such that its highest densities appear similar in both terminal and peri-centromeric regions, and the overall weight of Helitron elements is very concentrated in these regions compared to gene-rich areas of the chromosome arms (Fig. 7A).

**Table 7:**
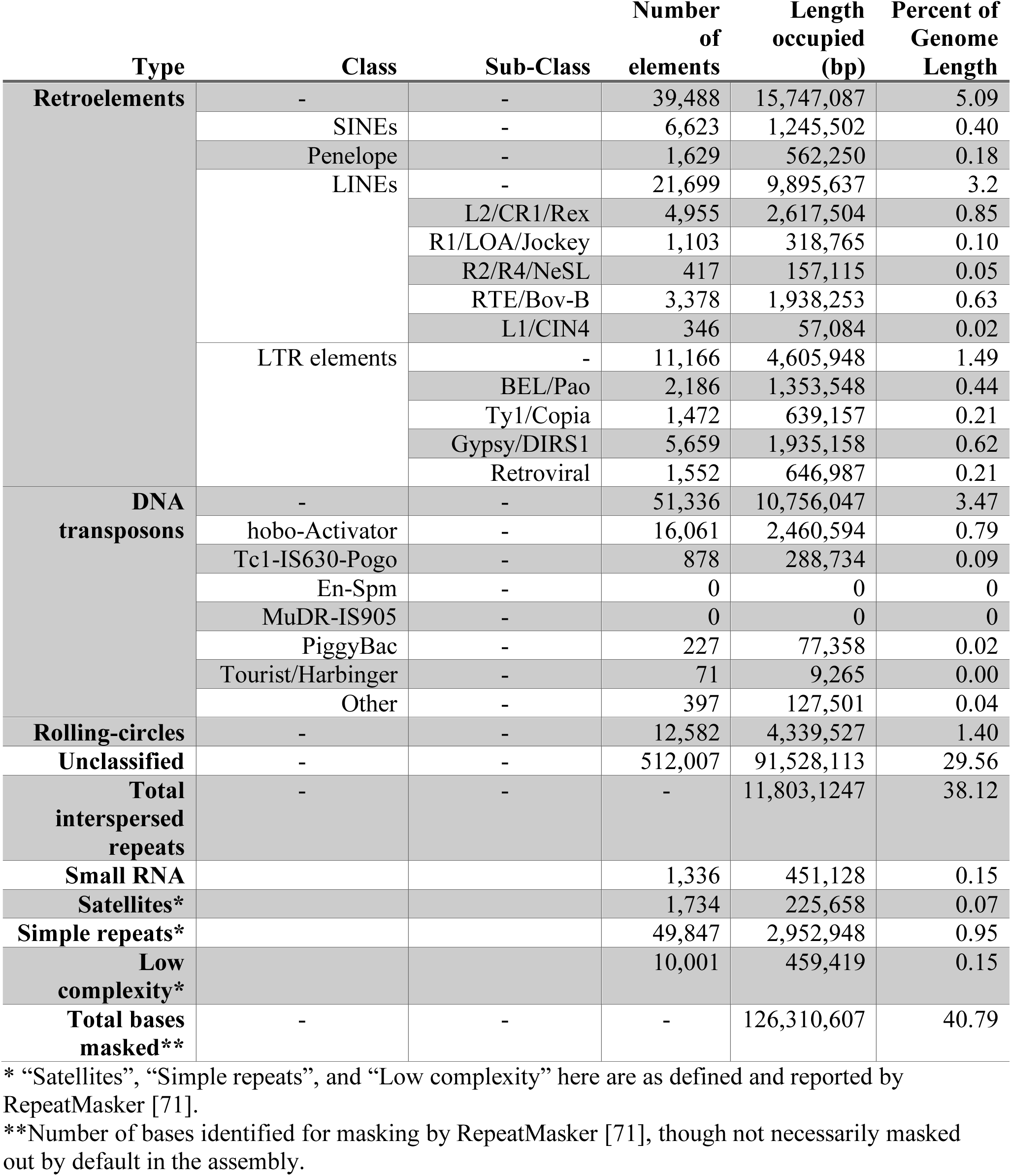
Repeat characterization of Bcop_v2 using RepeatMasker.

| Type | Class | Sub-Class | Number of elements | Length occupied (bp) | Percent of Genome Length |
| --- | --- | --- | --- | --- | --- |
| **Retroelements** | - | - | 39,488 | 15,747,087 | 5.09 |
|  | SINEs | - | 6,623 | 1,245,502 | 0.40 |
|  | Penelope | - | 1,629 | 562,250 | 0.18 |
|  | LINEs | - | 21,699 | 9,895,637 | 3.2 |
|  |  | L2/CR1/Rex | 4,955 | 2,617,504 | 0.85 |
|  |  | R1/LOA/Jockey | 1,103 | 318,765 | 0.10 |
|  |  | R2/R4/NeSL | 417 | 157,115 | 0.05 |
|  |  | RTE/Bov-B | 3,378 | 1,938,253 | 0.63 |
|  |  | L1/CIN4 | 346 | 57,084 | 0.02 |
|  | LTR elements | - | 11,166 | 4,605,948 | 1.49 |
|  |  | BEL/Pao | 2,186 | 1,353,548 | 0.44 |
|  |  | Ty1/Copia | 1,472 | 639,157 | 0.21 |
|  |  | Gypsy/DIRS1 | 5,659 | 1,935,158 | 0.62 |
|  |  | Retroviral | 1,552 | 646,987 | 0.21 |
| **DNA transposons** | - | - | 51,336 | 10,756,047 | 3.47 |
|  | hobo-Activator | - | 16,061 | 2,460,594 | 0.79 |
|  | Tc1-IS630-Pogo | - | 878 | 288,734 | 0.09 |
|  | En-Spm | - | 0 | 0 | 0 |
|  | MuDR-IS905 | - | 0 | 0 | 0 |
|  | PiggyBac | - | 227 | 77,358 | 0.02 |
|  | Tourist/Harbinger | - | 71 | 9,265 | 0.00 |
|  | Other | - | 397 | 127,501 | 0.04 |
| **Rolling-circles** | - | - | 12,582 | 4,339,527 | 1.40 |
| **Unclassified** | - | - | 512,007 | 91,528,113 | 29.56 |
| **Total interspersed repeats** | - | - | - | 11,803,1247 | 38.12 |
| **Small RNA** |  |  | 1,336 | 451,128 | 0.15 |
| **Satellites\*** |  |  | 1,734 | 225,658 | 0.07 |
| **Simple repeats\*** |  |  | 49,847 | 2,952,948 | 0.95 |
| **Low complexity\*** |  |  | 10,001 | 459,419 | 0.15 |
| **Total bases masked\*\*** | - | - | - | 126,310,607 | 40.79 |

\* “Satellites”, “Simple repeats”, and “Low complexity” here are as defined and reported by RepeatMasker [71].
\*\*Number of bases identified for masking by RepeatMasker [71], though not necessarily masked out by default in the assembly.

Repeats that could be recognized and classified as belonging to a particular repeat or transposon family tend to be enriched with proximity to centromeres and termini (Fig. 7A). This is also true for locations that match conserved classified repeats from across other arthropods (Fig. 7A). Thus, the regions associated with termini and around centromeres are enriched for repeats that are conserved enough to classify into families and/or be most similar to repeats from other species. However, the majority of repeat families in the repeat library trained *de novo* on the *B. coprophila* genome could not be classified according to known repeat families in the same automated fashion by the repeat modeling and annotation programs (Table 7), and are referred to as ‘unclassified repeats’. While the density of unclassified repeats is still moderately higher near centromeres and telomeres, the bias is much lower, and the unclassified repeats are much more spread out across the rest of the chromosome arms (Fig. 7A). It is possible some of these unclassified repeat families actually correspond to gene families or repeat families, newly arisen or otherwise, that are just not included as possible classification terms in the programs used. Unclassified repeats may also consist of highly degraded copies of known repeat families that were simply not recognized, which would indicate that repeat sequences are either better maintained near centromeres and near chromosome ends, or younger in those regions.

### Chromosome-scale scaffold termini highlight arrays of retrotransposon-related and satellite sequences as candidate telomeric repeats

The telomeres of most studied eukaryotes are composed of very short (5-10 bp) tandem repeats that are maintained by telomerase [15,16,47,48], most often similar to the mammalian hexamer (TTAGGG). For example, most insects have the pentamer, TTAGG [16]. However, most or all Dipterans studied have lost both the short tandem repeats and the telomerase gene, TERT [16]. The telomeres of *Drosophila*, a “higher” Dipteran, are composed of telomere-specific retrotransposons (TSRTs) that point towards the centromere and are maintained by their reverse transcriptase genes [15,16,47,48]. The telomeres of lower Dipterans are less well understood. *Rhynchosciara americana* appears to have 14-22 bp long tandem repeats, which might be maintained by a TSRT related to Drosophila TSRTs [16]. Non-biting midges (Chironomidae) have “Long Complex Terminal Tandem Repeats” (LCTTRs) with a unit length of 176 bp in one species and ∼350 bp in another species, although the latter appears to have evolved from a simpler 175 bp repeat [15,16]. The long repeats in midges might be maintained by homologous recombination or gene conversion, although long complementary RNAs have been observed, suggesting an RNA intermediate as with telomerase and TSRTs [15,16]. A mosquito has even longer repeat lengths of ∼820 bp in its telomeres, and at least one higher Dipteran, *Drosophila tristis*, has LCTTRs of similar length to midges (181 bp) [15]. In at least some midges, and in some insects more generally, telomeres can differ from each other, and some telomeric repeats may also be seen in centromeric regions [15,16,47,48]. We analyzed the updated *Bradysia coprophila* genome to explore the evidence for or against each telomeric system in this lower Dipteran. The chromosome-scale scaffolds allowed us to closely inspect the sequence at each terminus to hunt for candidate telomere repeat sequences, as has been done recently by others for many insects [47,48].

As expected for Dipterans, we found no evidence of the TERT gene in the *Bradysia coprophila* genome or proteome, and no evidence for short (5-10 bp) tandem repeats at the scaffold termini, including the canonical insect pentamer. We also did not find evidence for tandem repeats in the 14-22 bp range near the scaffold termini. What we did find at the scaffold termini were (i) arrays of retrotransposon-related sequences with unit lengths of 1.7-3.3 kb, (ii) arrays of LCTTRs (or satellites) with unit lengths of 161-191 bp, and (iii) examples of shorter tandem repeats in the 51-56 bp range further into the sub-telomeres. These are briefly described below with the caveat that one or more of the scaffold termini may not represent the true telomere sequences.

Retrotransposon-related sequences (RTRS) were enriched near the left termini of autosomal scaffolds (Fig. 7C-F – Pink triangles) and both termini of the X scaffold (Fig. 7G-H – Purple triangles). The left termini of scaffolds corresponding to chromosomes II, III, and IV are composed of arrays of RTRS with unit lengths of 1.7-3.3 kb. Although the RTRS repeat units have different lengths, they share ∼1.2 kb of sequence, and all three were classified in the repeat annotation as non-LTR LINE elements related to RTE-BovB, CRE, and/or CR1. This is reminiscent of *Drosophila*, which have non-LTR elements for telomeres. However, the RTRS arrays found at the very ends of the left termini of II and III point away from the centromere, which differs from *Drosophila*. On the left terminus of II, there are ∼8 copies of a 1.7 kb RTRS spanning the first ∼14 kb, and on III, there are ∼3-4 copies of a 3.3 kb RTRS spanning the first ∼12 kb (Fig. 7D-E). Each unit in these arrays has a single location with BLAST hits to reverse transcriptase (RT) proteins. After the RTRS array on III, the repeat annotation shows a centromere-facing LINE (RTE-ORTE). The left terminus of IV has a single left-facing LINE-related sequence and corresponding BLAST hit to RT proteins ∼10 kb into the scaffold. This is then followed by 13-14 copies of a centromere-facing 2.5 kb RTRS unit, although this unit does not have the region corresponding to RT protein BLAST hits despite being related to the RTRS units on II and III that do (Fig. 7F). In contrast to the left termini of the autosomal scaffolds, the right termini do not have RTRS arrays, at least not so close to the termini (Fig. 7I). The right terminus of II has 1-2 partial hits to the RTRS units in the sub-telomeres. The right terminus of III has two related hits: the very final ∼100 bp matches the RTRS units, and there is a centromere-facing RT protein hit to a region labeled as LINE/CR1 ∼10 kb from the end. Finally, ∼90-115 kb from the right terminus of IV are two sub-telomeric arrays of 2 and 6 centromere-facing RTRS units highly similar to those at the left terminus of IV. Of note to *Sciara* researchers, the 1.7-3.3 kb RTRS units do not appear to be related to the very abundant non-LTR ScRTE sequence and its corresponding F4 probe studied by Escribá et al [45], which were not found within 10 kb of any terminus, typically having a first instance 60-440 kb away from termini, much farther inward than the RTRS arrays described above. The left and right termini of the scaffold corresponding to the X chromosome also had retrotransposon-related sequence, but different from that on the autosomes. Both sides of the X scaffold start with sequence classified as a centromere-facing LTR/Pao elements, and both are followed immediately by terminus-facing DNA transposons (Fig. 7G-H). LTR elements would be surprising candidates for telomere sequences compared to *Drosophila*, which depends on the mechanism associated with non-LTR elements for telomere maintenance [16], and may indicate that the X telomeres are not well-represented on this scaffold.

Long Complex Terminal Tandem Repeats (LCTTRs or satellite repeats) were also enriched near the right termini of autosomal scaffolds (Fig. 7I; Table 8). Specifically, the right termini of scaffolds corresponding to chromosomes II and III end with ∼28 copies of a 175 bp satellite (Fig. 7I – Orange triangles; Table 8) and ∼28 copies of a 161 bp satellite (Fig. 7I – Red triangles; Table 8), respectively. The most striking example is the right terminal ∼40.3 kb of IV, which contains 229 copies of a 176 bp satellite (Fig. 7I – Blue triangles; Table 8). Each satellite species is very close to the LCTTR unit length seen in midges of 176 bp. Although each right terminus ends with a different satellite, each terminus or sub-telomeric region also highlights one or more of the others and/or of two other prevalent satellites with unit lengths of 191 bp and 56 bp (Fig. 7I – Brown and light purple triangles, respectively; Table 8). For example, the right terminus of II also has an array of the 191 bp (brown) satellite, the right-terminal sub-telomeric regions of all chromosome scaffolds have arrays of the 56 bp (purple) satellite, and arrays of the 161 bp (red) satellite are found in the right-side sub-telomeric regions of scaffolds for X, II, and IV. The scaffold for chromosome IV shows the richest examples of satellite arrays in its sub-telomeric regions, both the left and right termini of which have arrays of four or more of these satellite species. The left side of IV in particular is super-enriched for the 175 bp (orange) satellite that was seen at the right terminus of II. Interestingly, all RTRS units above have 1-4 partial hits to the 176 bp (blue) satellite sequence that was found in a long array concluding the right terminus of IV.

**Table 8:**
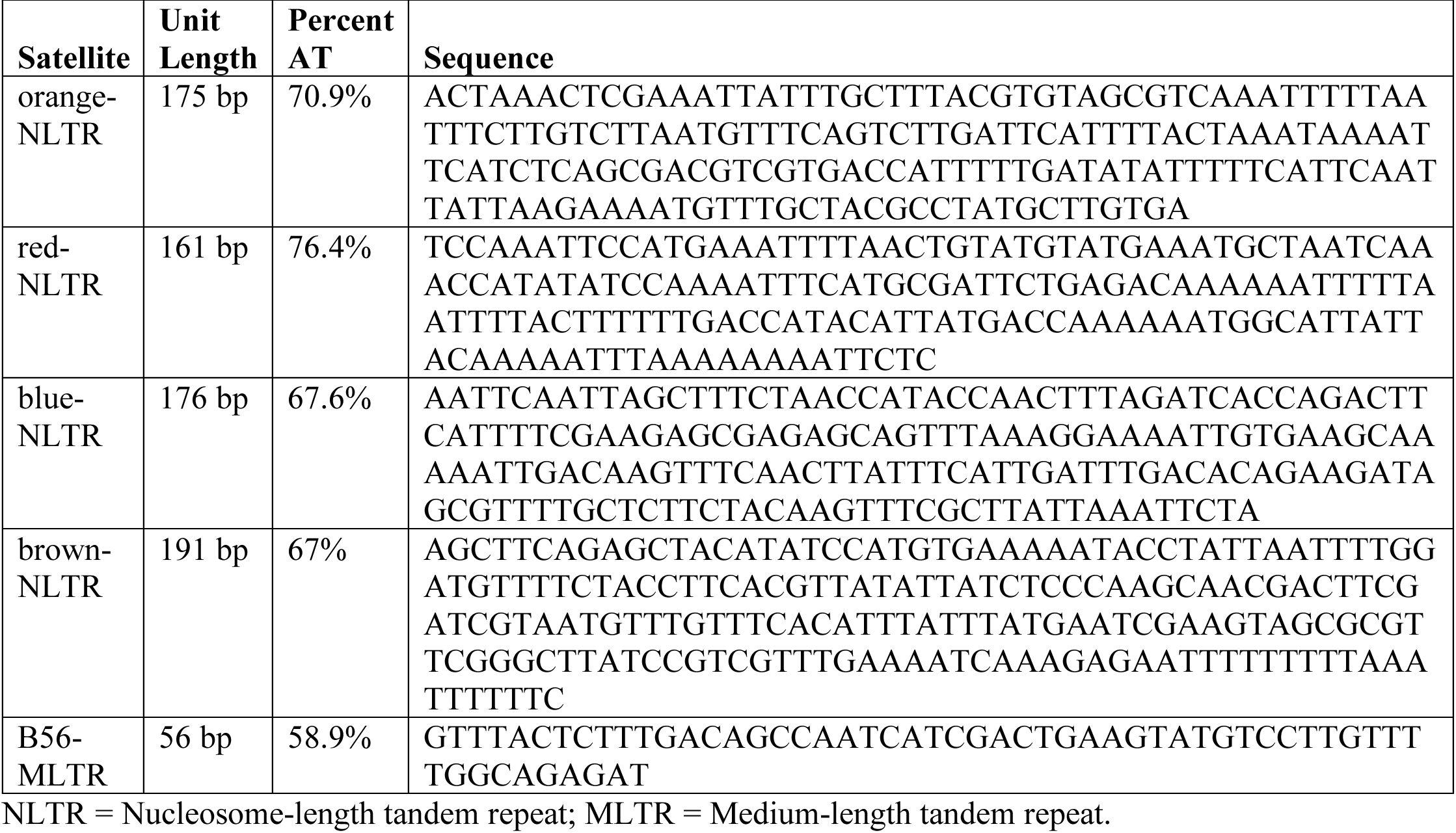
Sequence of satellites found at or near scaffold termini.

Centromeric and pericentromeric regions on the chromosome-scale scaffolds are also enriched for the termini-associated satellites (Fig. 7J). For example, the 161 bp (red) satellite was found in the centromeric regions of all three acrocentric chromosome scaffolds: X, II, and III. In fact, the 161 bp (red) satellite was tightly associated with arrays of the 143-155 bp centromeric sequence, Sccr (Fig. 7J, Yellow triangles), on the X scaffold, and also turned out to be part of the Sccr-linked repeat family that helped identify centromeric positions in scaffolds, especially for chromosome II (Fig. 7A; “Sccr family”). The centromeric region of II did not have strong representation from Sccr (likely to be an assembly artifact), but had 47 copies of the Sccr-associated 161 bp (red) satellite and multiple arrays of the 175 bp (orange) and 176 bp (blue) satellites. The only metacentric scaffold, IV, had massive arrays of Sccr, but was depleted for termini-associated satellites, although it had 131 tandem copies of the 191 bp (brown) satellite in the pericentromeric region ∼75 kb away from the closest Sccr repeats.

### Known biological features of the X chromosome further confirm chromosome identity and structural accuracy

There are three regions along the X chromosome where it folds back on itself such that these loci physically interact, leaving the X looking like a loop with its ends knotted up together on polytene spreads [22,23,30,31,49]. For convenience, Figure 8 shows examples of the X polytene chromosome doing this (Fig. 8A-B), but also see previous works [22,23,31,45,49]. These regions have been referred to as “repeats on the X” [31] or “repeat regions R1, R2, and R3” [22,49]. The richest descriptions of them simply refer to their banding pattern when polytenized as “three three-band non-adjacent pattern repeats” [22]. However, whether these regions truly are repeats of each other at the level of DNA sequence or whether they even contain repetitive DNA sequences is yet unknown. Therefore, we refer to the three “repeat” loci on the X here simply as the “Fold-Back Regions” (FBRs 1-3). Each locus is reported to be composed of three polytene bands, which indicates they correspond to DNA lengths of tens to hundreds of kb, assuming similar band lengths as *Drosophila* [50]. The Fold-Back Regions have been mapped cytogenetically to the zones within the X polytene map (Fig. 8C), and are all downstream from the centromere, with FBR1 being closest and FBR3 being furthest away [22,23,30,49]. Specifically, slightly downstream of the centromere, which is part of locus X/1A, is FBR1 at locus X/1C. FBR1 is followed within a relatively short distance by FBR2 at locus X/4A. A relatively longer distance separates FBR2 from FBR3 at locus X/12C, which is closest to the distal (non-centromeric) end of the chromosome that ends at locus X/14C (Fig. 8C).

**Figure 8:**
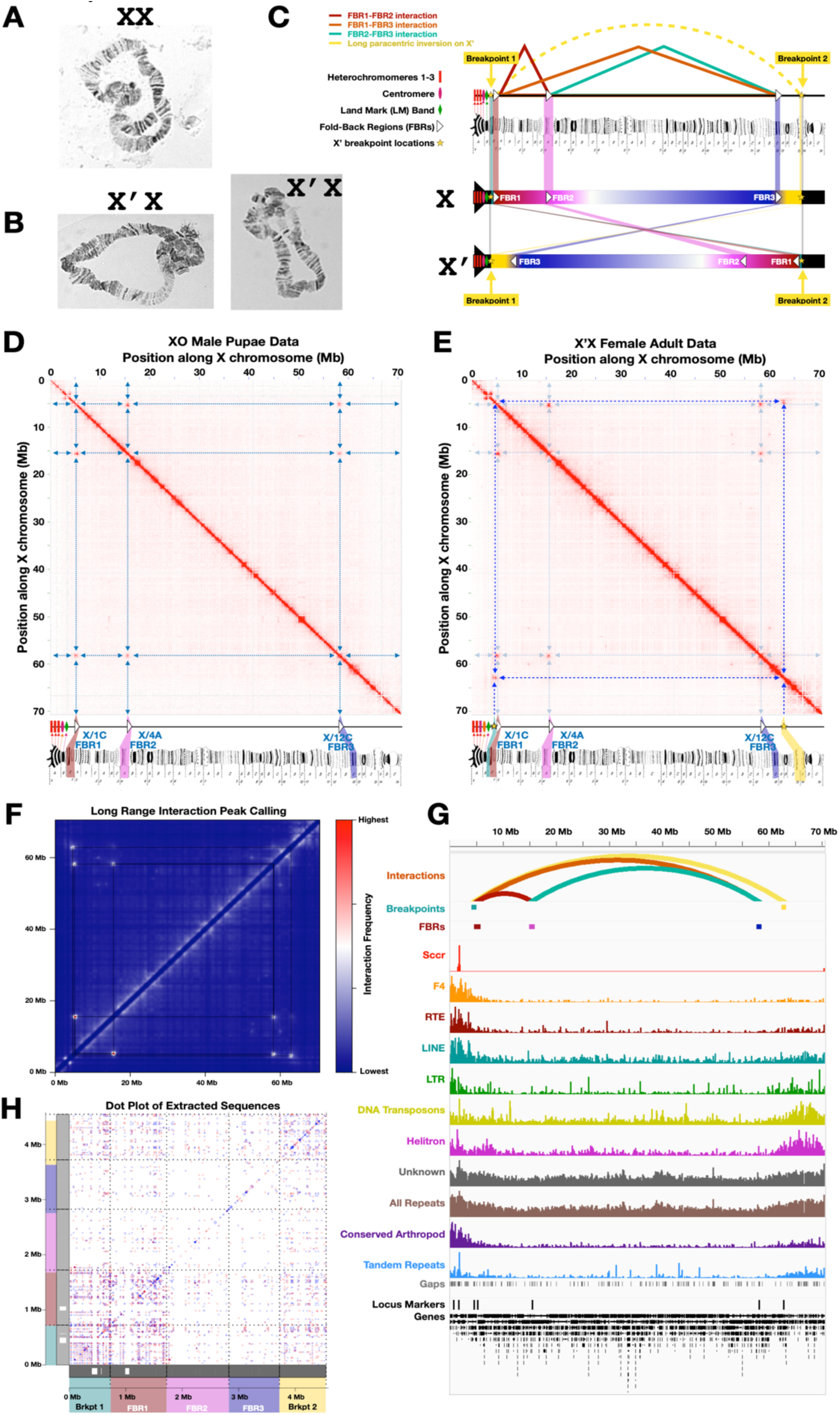
Putative Fold-Back Regions (FBRs) and X’ Breakpoints further confirm accuracy of the X scaffold. (A-B) Polytene chromosome spreads demonstrating how the X and X’ chromosomes are usually found: folded back on itself with specific loci physically interacting. **(C)** Pictorial representation of what is known about the X and X’ chromosomes, FBRs, and inversion breakpoints. **(D)** Hi-C map from male pupae data showing three interacting loci and how they appear to correspond to the expected locations of FBRs along polytene maps. **(E)** Hi-C map from X’X adult female data [10] that shows the FBRs as well as two additional loci that correspond to the long paracentric inversion breakpoints. **(F)** An intermediate scatter plot heat map visualizing part of the long-range interaction peak cluster calling pipeline with squares demonstrating long-range interactions found by this method. **(G)** IGV Trace of the X chromosome with special attention to the locations identified as FBRs and breakpoints (top 3 tracks), as defined by our long-range Hi-C interaction peak detection method, and how they relate to repeat-rich and gene-rich regions. **(H)** Dotplot of extracted sequences corresponding to FBRs and breakpoint regions, showing sequence matches within and between each sequence. Blue dots represent matches on the same strand. Red dots represent matches on opposite strands. Polytene chromosome maps in C-E were reproduced from plates 1, 2 and 3 of Gabrusewycz-Garcia (1964) [30] with permission from Springer Nature under permission number: 5490830617309.

There is a variant of the X chromosome called the X’ that is found only in female-producing females [1,2,10,22,49,51–54], which also folds back on itself via the fold-back region interactions. The X’ has long been known to differ from the X by a long paracentric inversion that reverses the order of the fold-back regions such that FBR3 is centromere-proximal and FBR1 is most distal (Fig. 8C). An example X’X polytene folding back on itself via FBR interactions is specifically shown in Fig. 8B; note the lack of synapsis between the chromatids from different homologs (X and X’) across most of chromosome length corresponding to the long inversion compared to the XX polytene in Fig. 8A. The breakpoints that correspond to the long paracentric inversion have been mapped cytogenetically on the polytene zones upstream of FBR1 and downstream of FBR3 (Fig. 8C) [22,49], and more recently have been explored bioinformatically using sequencing data from X’X females and the Bcop_v2 X chromosome scaffold reported here [10]. Using images of the X’ breakpoints and FBRs [22] with polytene maps [30], we estimate that (i) the 5’ breakpoint is downstream of both the centromere and the so-called dark “landmark” band [22,45] in an interband quite close to FBR1 (X/1C), likely at the end of X/1B or the beginning of X/1C, and (ii) the 3’ breakpoint occurs downstream of FBR3 (X/12C), likely at the end of X/13C or beginning of X/14A (Fig. 8C).

If the Fold-Back Regions physically interact in the X chromosomes of male pupae (for which we collected Hi-C data), then they should appear on the Hi-C map of the X chromosome scaffold as three off-diagonal dots with high frequency long-range interactions. Indeed, the Hi-C map of the X chromosome shows dots corresponding to the frequent interactions of three distal loci in three-dimensional space (Fig. 8D). Moreover, Hi-C data from X’X adult females [10] shows the same three high frequency long-range interactions along the X scaffold, but also has two additional apparently-long-distance interactions corresponding to the breakpoints of the long inversion (Fig. 8E). Importantly, the locations of and nucleotide distances between the centromere, FBRs 1-3, and the X’ breakpoints follow expectations based on the polytene maps (Fig. 8D-E, Table 9, see Methods). FBR1 is expected to be within ∼3.4 Mb (∼2 sub-zones) of the centromere and up to ∼839 kb (∼1/2 sub-zone) downstream of the centromere-proximal X’ breakpoint. Indeed, putative FBR1 is ∼3.49 Mb and ∼620 kb downstream from the centromere and inversion breakpoint, respectively (Table 9). Moreover, FBR1 and FBR2 are expected to be separated by ∼11.8 Mb (∼7 sub-zones), and FBR2 and FBR3 are expected to be separated by 43.6 Mb (∼26 sub-zones). The observed distances are 10.3 Mb and 42.7 Mb, respectively (Table 9). The 3’ breakpoint is expected to be ∼5 Mb downstream (∼3 sub-zones) of FBR3, and was found to be 4.65 Mb downstream. Finally, FBR3 was predicted to be ∼10.1 Mb upstream of the distal end of chromosome X, and the observed scaffold location is 12.3 Mb upstream (Table 9). In all cases, the observed distances and locations were exceptionally close to projections based on known polytene map distances and locations. Thus, it is extremely likely that the three distal loci that physically interact at high frequencies according to Hi-C data are indeed the three Fold-Back Regions, which have not been previously known at the sequence level. The Hi-C interactions in male pupae and female adults indicate that the physical interactions of the X chromosome seen in polytene spreads are not *in vitro* artifacts, nor are the interactions limited to polytenes in larval salivary glands, as they now have been observed *in vivo* across different tissues and life stages in both sexes (male pupae and female adults). Overall, the FBR and X’ breakpoint locations further confirm the identity and structural accuracy of the X chromosome scaffold.

**Table 9:** Nucleotide distances between FBRs and other features of the X chromosome are concordant with expectations. The X polytene map [30] is organized into 14 zones of similar lengths, each with 3 subzones (A, B, and C) for a total of 42 sub-zones. Assuming nucleotide lengths are also similar, the ∼70.5 Mb X-scaffold would indicate zone lengths of ∼5 Mb (5.036276) and an average sub-zone length of ∼1.68 Mb (1.678759), which we used to compute expected nucleotide distances.

| **Locus 1** | **Locus 2** | **Approx. Polytene**<br>**Map Distance (#**<br>**sub-zones)** | **Expected**<br>**Distance (Mb)** | **Observed**<br>**Distance (Mb)** |
| --- | --- | --- | --- | --- |
| Centromere | FBR1 | 2 | 3.4 | 3.49 |
| 5' X' break point | FBR1 | 0.5 | 0.839 | 0.620 |
| FBR1 | FBR2 | 7 | 11.8 | 10.3 |
| FBR2 | FBR3 | 26 | 43.6 | 42.7 |
| FBR3 | 3' X' break point | 3 | 5.0 | 4.65 |
| FBR3 | 3' end of<br>chromosome | 6 | 10.1 | 12.3 |

The three Fold-Back Regions on the X chromosome have been described as repeats in cytogenetic studies [22,23,31,49]. However, the sequences were previously not known, and whether they are composed of repeats has not been well-established. Given the likeliness that the three loci on the X with long-range Hi-C interactions correspond to the three FBRs, their scaffold locations and sequences were extracted and analyzed. FBR scaffold locations were discovered by calling long-range Hi-C interaction peaks, after first filtering out short interactions less than 5 Mb (Fig. 8F-G; see Methods). The length of each FBR was in the ∼103 kb to ∼1.1 Mb range with interaction frequency summits in the 34-69 kb range, depending on how boundaries were defined (see Methods). To maximize the chance of detecting long stretches of similarity among these regions, we used the longest estimates. When directly comparing the extracted sequences of these three loci by BLAST, there were stretches of similarity found in all three (Fig. 8H). Although some matches reached the kb range, most stretches were short (< 200 bp), and most or all were associated with repeats found elsewhere in the genome. The FBRs did not have shared repeat families specific to these regions. Looking at the distribution of repeats across the X chromosome, it is plausible that the first locus, putative FBR1, is embedded in a highly repetitive region, but less likely for the two downstream loci (putative FBRs 2 and 3), which appear to be in relatively unique regions (Fig. 7A-B, Fig. 8G). When comparing the entire X chromosome to itself using dotplots, if these regions had high similarity the length of up to three polytene bands, there would be off-diagonal “dots” on the dotplots corresponding to the locus pairs analogous to that seen in the Hi-C interaction frequency plots. However, such long stretches of similarity are not seen in those off-diagonal locations (Fig. 7B). Overall, the evidence so far suggests that although the three FBR loci interact, it is not necessarily due to vast stretches of homology that would span 1-3 polytene bands (tens to hundreds of kb). Although these three interacting regions are not necessarily three repeated loci, this does not exclude the possibility that one or a few of the shorter stretches shared between the regions are important aspects of their physical interactions. In contrast to FBRs 2 and 3 being in relatively repeat-poor regions of the X chromosome, the X’ inversion breakpoints are both buried in repetitive domains corresponding to peri-centromeric and sub-telomeric regions, which do share more similarity with each other (Fig. 7A-B, Fig. 8G). The 5’ breakpoint and nearby putative FBR1 are embedded in a repeat block seen on the X scaffold dotplot near the centromeric (left) end of the chromosome that has high similarity with the right terminal repeat block (Fig. 7A-B, Fig. 8G). The presence of the FBRs on both the X and X’ suggest the FBRs and their interactions preceded the inversion, and it is possible that the evolution of long paracentric inversion on the X’ was mediated by the systematic fold-back interactions of FBR1 and FBR3 bringing these peri-centromeric and sub-telomeric repeat blocks close together.

In addition to repeats, there are also genes within the boundaries of all three putative FBR loci (as well as for the X’ inversion breakpoints). Thus, it is also possible that the long-range fold-back interactions may be involved in regulation of a gene or genes in the fold-back regions, or that these regions each have representatives from a shared gene family. To maximize the sensitivity of finding a gene family with members in all three regions, we focused again on the broader region boundaries, not the narrowest estimates. Depending on the gene annotation set and FBR boundaries used, there are 85-203 genes across all three FBRs with 35-78 in FBR1, 29-80 in FBR2, and 21-45 in FBR3. Regarding whether there are homologous genes in these regions, some of the stretches of nucleotide similarity shared by all three FBRs do overlap gene boundaries, but almost exclusively in introns, with very few exon hits and no coding sequence (CDS) overlaps. In agreement, there appears to be no homologous protein family represented in all three regions when analyzing protein sequence similarity with BLASTP. The regions of similarity at the nucleotide level within introns are largely from transposons and other repeats. Nevertheless, we cannot exclude the possibility that some stretches of similarity in the introns or near genes in general are regulatory elements shared by the FBRs. It is also possible that FBR-specific sequences, not found elsewhere in the genome, are important. There are millions of FBR-specific kmers of various sizes (15-80 bp) not found elsewhere in the genome. These kmers correspond to on the order 0.5-1 Mb of sequence from hundreds to thousands of sub-regions of the FBRs up to hundreds of bp in length, which also lack BLAST hits elsewhere in the genome. Approximately 70% of this sequence is within genes and a partially-overlapping ∼16-17% corresponds to highly diverged repeats, but ∼23.5% is intergenic and not labeled as a repeat. Thus, there may be intergenic (and genic) FBR-specific nucleotide elements that are found in one or more of the FBRs, but not elsewhere in the genome. Overall, it will be exciting to further elucidate in future studies what sequences and proteins mediate the long-range fold-back interactions, what function the long-range interactions serve, whether these regions and interactions are involved in other aspects of X chromosome biology, and if these interactions play a role in facilitating structural rearrangements on the X and X’ chromosomes.

## Discussion

*Bradysia coprophila*, a dark-winged fungus gnat from the lower Diptera (Nematocera), has interesting chromosomal biology that includes developmentally-programmed examples of elimination, non-disjunction, polytenization, intrachromosomal amplification, and maternally-acting sex chromosomes [1,2]. It also presents opportunities for studying sex chromosome evolution [10,55] and horizontal gene transfer [6–8]. A crucial part of the toolbox for studying these phenomena is a genome sequence. The previous reference genome sequence for *Bradysia coprophila*, Bcop_v1, was a high quality, high contiguity assembly of the somatic genome (chromosomes X, II, III, IV) produced from PacBio RS II and Oxford Nanopore MinION long read data from male embryos, and subsequently scaffolded with optical maps of male pupae DNA molecules from BioNano Genomics [9]. Bcop_v1 was produced through an extensive evaluation process comparing nearly 100 assemblies [9]. Moreover, as *Bradysia coprophila* males have only one copy of the X chromosome, but two copies of autosomes, all contigs in Bcop_v1 were classified as X-linked or autosomal [9]. Finally, two gene annotation sets were constructed for Bcop_v1: we produced one using Maker2 [9,33,34] and NCBI produced one for RefSeq [9,35]. However, Bcop_v1 was not chromosome-scale and mostly not anchored into chromosomes, limiting its usefulness in studying the evolution and behaviors of *Bradysia coprophila* chromosomes.

Our goal in this project was to improve the reference genome assembly for *Bradysia coprophila* to better-facilitate studies of its interesting chromosome behaviors and evolution. To do so, we upgraded Bcop_v1 [9] to chromosome-scale scaffolds (Bcop_v2) using Hi-C technology. The updated assembly of the *Bradysia coprophila* somatic genome (Bcop_v2) represents each chromosome (X, II, III, IV) as its own scaffold for the first time, and is oriented in the same direction as polytene maps produced in the 1960s [30] to be consistent with historical research. The chromosomal identities of the scaffolds were established and supported several ways, including multiple anchor sequences, chromosome lengths, depth of coverage, centromere positions, and other idiosyncrasies of the X chromosome. The accuracy of the chromosomal structure of the scaffolds was also supported several ways, including the concordance of anchor sequence locations along the scaffolds with their locations along polytene maps, the relative positioning of centromeres and telomeres, the increasing enrichment of repeats with proximity to centromeric and telomeric regions, and other landmarks on the X chromosome (FBRs and X’ breakpoints). As studies are ongoing across multiple research groups with the gene annotation sets produced for Bcop_v1, we lifted over these gene annotation sets directly onto the chromosome-scale scaffolds so researchers can seamlessly transition ongoing studies from Bcop_v1 to Bcop_v2. The chromosomal specificity of the genes within each chromosome scaffold was further supported by comparative genomics, which showed that the chromosomes of more closely related species were far more similar in terms of gene sets than more distantly related species, both visually and using entropy statistics, the latter of which also reproduces expected phylogenetic species groupings. The totality of evidence overwhelmingly suggests that Bcop_v2 contains high quality scaffolds of the four somatic *B. coprophila* chromosomes (X, II, III, IV).

The rich history of studying *B. coprophila* chromosomes cytogenetically with respect to polytene map zones [30] has been a major benefit in producing and interpreting these chromosome-scale scaffolds. For example, previous *in situ* hybridization studies that identified the chromosomal addresses of cloned DNA sequences [17–29] has allowed us to identify the chromosomal identity of each chromosome-scale scaffold, orient them according to the polytene maps, test their structural accuracy, map their centromeric regions, and even to study known features of the X corresponding Fold-Back Regions and breakpoints of a long paracentric inversion on a variant of the X chromosome called the X’ (X prime). Translating the rich history of cytogenetic observations could be aided by more extensively annotating the polytene map zones within the Bcop_v2 chromosome-scaffold sequences. In the future, the polytene zones can roughly be mapped using a combination of landmark sequences identified by FISH, genomic profiling techniques, and assumptions about the sizes of the polytene zones. It may also be possible to re-purpose the Hi-C data reported here to identify the boundaries of bands and interbands seen in polytenes, which would help interpret the boundaries of the historical polytene zones and sub-zones. Polytene bands and interbands have been associated with topologically associated domains (TADs) in *Drosophila* polytene chromosomes using Hi-C data [56,57]. The locations and sequences of the polytene zones could help re-interpret older cytogenetic studies through a genomic lens. Regardless, the male (XO) pupae Hi-C paired-end reads will continue to be useful for correcting mis-assemblies and scaffolding future *Bradysia coprophila* genome assemblies produced with newer long-read datasets (e.g. PacBio HiFi or ultra-long Q20 nanopore reads).

As Bcop_v2 is derived from scaffolding Bcop_v1, there were no significant changes to conclusions about the types and amounts of repeats reported previously [9]. However, the repeats in Bcop_v1 are scattered across hundreds of unanchored contig sequences whereas the scaffolds of Bcop_v2 allow us to see where the repeats are located within the chromosomes. We found recognizable repeat families concentrated around peri-centromeric and sub-telomeric regions with different propensities. Retrotransposon families in general, and especially those classified as LINE (non-LTR), were very enriched in pericentromeric regions, including the LINE/RTE sub-family as shown to be enriched around centromeres previously [45]. Despite a major bias towards centromeric regions, retrotransposon related sequences and reverse transcriptase BLAST hits are found throughout scaffold terminal regions as well, including within 10 kb of most termini. Retroelements have also been shown to be enriched in functional *Drosophila* centromeres [58], and to make up and play an important role in maintaining *Drosophila* telomeres [15,16]. Helitron elements were enriched in both terminal and centromeric regions. DNA transposon families were biased more towards scaffold terminal regions, although they are likely sub-telomeric as they do not appear as terminal sequences. Ultimately, defining the repeats that make up *B. coprophila* telomeres and how those telomeric repeats are maintained is beyond the scope of this paper. However, we checked for signatures of known telomeric systems. As with all Diptera, Telomerase Reverse Transcriptase (TERT) was not found in the *B. coprophila* genome sequence nor proteome, nor was the associated hexamer/pentamer (TTAGGG/TTAGG) or other variant found at scaffold ends. Non-LTR retrotransposons (HeT-A, TART, TAHRE) make up and maintain Drosophila telomeres [15,16], and TART copies were also found near the chromosome ends of more closely related species, *Rhynchosciara americana* [16]. Although all these transposons are possibly too far diverged to find matches in the terminal regions of *B. coprophila* chromosome scaffolds (Bcop_v2), reverse transcriptase genes can certainly be found in the sub-telomeric regions, even within the first 10 kb. Moreover, the left termini of the *B. coprophila* autosomal scaffolds all seem to have arrays of retrotransposon-related sequences (RTRS) with units in the 1.7-3.3 kb range and both termini of the X scaffold were associated with retrotransposons in the repeat annotation. *Rhynchosciara* was also shown to have a “special type of short tandem repeats” (stSTRs) of 14-22 bp [16]. Given their close evolutionary relationship, *B. coprophila* might be expected to have the same system. However, we did not find short repeats at the *B. coprophila* chromosome scaffold termini, although tandem arrays of short repeats may be found further inward in the sub-telomeric regions. It is also possible *B. coprophila* has Long Complex Terminal Tandem Repeats (LCTTRs) like those found in Chironomidae and Culicidae [15,16]. Indeed, we found a set of satellite sequences with unit lengths of 161-176 bp associated with the right termini of autosomal scaffolds. These terminal and sub-terminal satellites were also seen in peri-centromeric regions as was also seen in some midges [15]. We note that the repeat lengths of these satellites are similar to nucleosome spacing lengths. For example, the length of DNA per nucleosome in Drosophila is ∼175 bp [59], which includes the linker region, and two of these candidate repeats are exactly 175-176 bp. Interestingly, all 1.7-3.3 kb RTRS units above have 1-4 partial hits to the 176 bp satellite sequence, suggesting an ancient relationship between these satellites and transposons. This is also reminiscent of the cycling between retroelements and satellite DNAs found in the centromeres and telomeres across several *Drosophila* species [60]. Regardless, our current model for *B. coprophila* telomeres based on the termini of these chromosome-scale scaffolds would appear to need to include arrays of both RTRS and LCTTR satellites. It is possible that the centromere-proximal (left) telomeres follow a different model than their distal (right) counterparts. In these scaffolds it appears that autosomal centromere-proximal termini, or all autosomal left termini, are correlated with RTRS arrays whereas the centromere-distal (or right side) termini are more correlated with LCTTRs. In insects, different telomere sequences have been found on different chromosomes, and even on the opposite sides of the same chromosome [15,16,47,48]. However, one or more of the *B. coprophila* scaffold termini may not be representative of the true telomeric sequence, and a definitive conclusion cannot yet be made for specific telomeres. The X scaffold in particular needs further scrutiny and refinement as so much of the intriguing chromosome biology in *B. coprophila* involves the short arm of the X chromosome specifically. In addition to experimental tests, such as in situ hybridization, newer PacBio HiFi and Q20+ ultra-long nanopore data will also shed more light on the telomere sequences in future work.

The chromosome-level nature of Bcop_v2 will allow researchers to design experiments and analyze data in a chromosome-specific way. The location and sequences of the telomeres and centromeres are now more obvious and open to future investigation. The X chromosome scaffold will prove useful in identifying the Controlling Element, an rDNA-associated sequence embedded in the short arm of the X upstream of the centromere. Moreover, the X chromosome scaffold has already benefited studies of the X chromosome variant, X-prime (X’), which differs from the X by a large paracentric inversion [10]. There are also interesting biological signals in the Hi-C maps that can be explored further. The notable example here are the three dots on the X chromosome Hi-C map that appear in the expected pattern of the three fold-back regions (FBRs) identified by Crouse [22,23,31,49]. Now that the fold-back region sequences have been identified within the scaffolds, it will be possible to begin defining what mediates their interactions and what function is served by doing so. Finally, the *B. coprophila* chromosome scaffolds will allow proper studies of synteny and chromosome evolution across closely and distantly related flies, and may prove useful for researchers working with closely related species if they are working with fragmented genome assemblies by letting them cluster the contigs into likely chromosome groups, and even ordering and orienting them into pseudo-chromosome scaffolds.

## Methods

### Male Pupae Collection

The dark-winged fungus gnat, *Bradysia coprophila*, was previously referred to as *Sciara coprophila* in chromosomal and molecular biology research papers since the 1920s [1,2,61], and is also known by other names such as *Sciara tilicola* and *Sciara amoena* [2]. In this study, the fungus gnats were from the HoLo2 line maintained in the International *Sciara* Stock Center at Brown University (https://sites.brown.edu/sciara/). *B. coprophila* females are monogenic, meaning they have either only male or only female offspring, which is determined by whether they harbor a variant of the X chromosome (X’) or not. Specifically, X’X females are female producers and XX females are male producers. The Holo2 line has a phenotypic marker gene on the X’ chromosome called *Wavy* [1,53]. Female producers (X’X) have the *Wavy* wing phenotype whereas male producers (XX) have wild-type straight wings. To collect only male pupae, crosses between straight-winged females (XX) and males (XO) were used to obtain strictly male progeny. Specifically, late-stage larvae from male-only matings were transferred from mating vials to agar petri plates where food and debris were removed from them, and where they entered pupation asynchronously over 2-4 days. The mixed-stage pupae were transferred to Eppendorf tubes (90-100 per tube), washed twice with 2X SSC, and then were flash frozen in liquid nitrogen and stored at −80°C until needed. Approximately 1,000 frozen male pupae collected across 11 tubes were shipped to Phase Genomics (Seattle, WA) on dry ice where they used the material as needed to prepare a Phase Genomics Proximo Hi-C sequencing library.

### Hi-C Data Collection

The frozen male pupae were ground into a fine powder. Chromatin conformation capture data was then generated using a Phase Genomics (Seattle, WA) Proximo Hi-C 2.0 Kit, which is a commercially available version of the Hi-C protocol [11]. Following the manufacturer’s instructions, intact cells from two samples of finely ground frozen male pupae were crosslinked using a formaldehyde solution, digested using the Sau3A1 (^GATC) restriction enzyme, and proximity-ligated with biotinylated nucleotides. This creates chimeric molecules made from DNA fragments that came close together *in vivo*, but that may not be close together along the genome sequence. As instructed by the manufacturer’s protocol, the chimeric molecules were pulled down with streptavidin beads and processed into an Illumina-compatible sequencing library. Sequencing was performed on an Illumina NovaSeq, generating a total of 115.2 million 2×101 bp paired-end reads.

### Hi-C assembly correction

The input assembly to Hi-C correction was Bcop_v1 [9], a mega-base scale, optical-map-scaffolded long-read assembly. Although Bcop_v1 consists of both scaffolds (two or more contigs joined by optical maps) and singleton contigs, here and throughout, the term “contigs” is used generically for both types of sequences from Bcop_v1, especially in the context as input to Hi-C guided contig correction and scaffolding. The Hi-C data was processed, as described below, according to Phase Genomics recommendations [62]. To produce a corrected assembly, the reads were aligned to the 205 primary contigs of Bcop_v1 [9] using BWA-MEM [63,64] with the −5SP and -t 8 options specified, and all other options default. The output was piped into SAMBLASTER [65] to flag PCR duplicates (to later exclude from analyses) and subsequently piped into SAMtools [66] using “-F 2304” to remove non-primary and secondary alignments. Juicebox [67–69] was used to manually identify and break contigs at putative mis-joined regions based on disruptions in the expected pattern from Hi-C alignments along the contigs [11,67–69]. Such putatively mis-joined regions were cut out and marked as “debris”. The cut-out debris ranged from ∼1.5-50 kb. Regarding the number of mis-joins detected and corrected (see Results), we leaned toward over-correction rather than under-correction at this step since the Phase Genomics Proximo Hi-C scaffolding in the next step can “repair” these breaks by joining the sub-contigs back together if breaking them apart was in error.

### Hi-C assembly scaffolding

The paired-end Hi-C reads were then aligned to Bcop_v1_corrected using the same alignment procedure as above, and these alignments were the input to the Phase Genomics’ Proximo Hi-C genome scaffolding platform, which was used to create chromosome-scale scaffolds from the Bcop_v1_corrected as described in Bickhart et al [13]. As in the LACHESIS method [14], the Proximo scaffolding process computes an interaction frequency matrix from the aligned Hi-C read pairs that is normalized by the number of DPNII restriction sites (^GATC) on each contig. Proximo then constructs scaffolds by optimizing the expected interaction frequencies and other statistical patterns in Hi-C data. Using a brute-force approach, approximately 20,000 separate Proximo runs were performed to optimize the number of scaffolds and the concordance with the observed Hi-C data. Finally, Juicebox [67] was again used to manually inspect for mis-join signals corresponding to scaffolding errors. The associated contigs from Bcop_v1 that were not included during the Hi-C scaffolding process were added back. Note that we also tried including the Bcop_v1 “associated contigs” (haplotigs, etc) during the Hi-C scaffolding process. In doing so, we found that although the resulting scaffolds were extremely similar to those produced from using only the primary contigs, it resulted in erroneously stitching together primary and associated contigs previously identified as haplotigs rather than keeping them separate. Therefore, the results of using only the primary assembly were preferred.

At the end of the scaffolding and correction process, there were 58 separate sequences not placed into the four chromosome-scale scaffolds (“unplaced”): 57 unscaffolded input contigs and a short (363 kb) Hi-C scaffold of 6 input contigs. Of the 57 unscaffolded input contigs, 46 were marked as “debris” (the regions cut out during the Hi-C correction step above). The “debris” was largely comprised of optical-map gap sequences from the input assembly. Six unscaffolded debris sequences ranging from ∼25-50 kb that had 100% N content were removed from the assembly (leaving 52 unplaced sequences). Furthermore, 24 unscaffolded sequences had leading or trailing NNNNN-content ranging from 1,489 - 36,461 bp, making up ∼6 - 96% of the unscaffolded sequences in which they resided, that was trimmed off. Only ∼19.5 kb of internal N content remained in the cleaned-up debris regions after removal and trimming of N-gap sequence. The primary assembly for Bcop_v2 consists of the four chromosome-scale scaffolds whereas the associated assembly for Bcop_v2 consists of the 52 cleaned-up unplaced sequences as well as all the “associated contigs” from Bcop_v1. For simplicity, even though the associated assembly of Bcop_v2 contains a scaffolded sequence, they are all referred to as ‘associated contigs’ throughout. In Bcop_v2, the retained BioNano Genomics optical map gap sequences are represented as capital N’s in the assembly and are of variable estimated lengths whereas the Hi-C gap sequences are represented as lower-case n’s and are always arbitrarily set to 100 bp in length.

### Anchoring and orienting the chromosome-scale scaffolds

Sequences with known chromosomal locations based on previous *in situ* hybridization work [17–29] [17–21,23–29] were used with BLAST [70] to identify the corresponding chromosome for each scaffold (Figures 2-3, Tables 2-3). For localization of centromeric and pericentromeric sequences within the chromosome-scale scaffolds, the short Sccr sequences [45] were used to identify Sccr-enriched Repeat Families in our custom *de novo* repeat libraries learned by RepeatModeler on the previous version of this genome (Bcop_v1) [9]. Sccr and Sccr-associated Repeat Families were mapped to Bcop_v2 using RepeatMasker [71]. To visualize the coverage and lengths of Sccr-associated alignments in IGV [72], and to identify the most likely centromere positions in the scaffolds, the lengths of all Sccr-repeat-family-associated alignments over a given position were summed for each position in the genome. The proportion of bases covered by Sccr-related or F4-related sequence in 100 kb bins was also computed with BEDtools [73] and visualized in IGV [72]. Approximated centromere positions within the scaffolds were used to compare to known centromere positions within the polytene chromosomes [22,30,31,45]. The anchor sequences and centromere positions were used to orient the chromosome-scale scaffold sequences to be concordant with conventional polytene maps [30]. Polytene map orientation was the final processing step for Bcop_v2, which was deposited to NCBI (GenBank GCA_014529535.2; WGS VSDI02; release date 2023-01-04). For researchers interested in how the contigs in Bcop_v1 correspond to Bcop_v2, we have also deposited AGP and BED files mapping these two assemblies together to: https://github.com/JohnUrban/Bcop_v2.

### Other Hi-C analyses on Bcop_v2

After correction, scaffolding, post-processing, cleaning, anchoring, polytene orientation, and otherwise all finishing steps to produce the final updated reference genome (Bcop_v2), the paired-end Hi-C reads were realigned to Bcop_v2 and interaction frequencies were computed and visualized similar to the above pipelines for correction and scaffolding: the output of mapping reads with BWA-MEM (−5SP -t 16; version 0.7.17-r1198-dirty) [63,64] was piped into SAMBLASTER (--removeDups; version 0.1.26) [65] to mark and remove duplicates, and further piped into SAMtools “view” (-F 2316; version 1.15.1 using htslib 1.15.1) [66] to filter out alignments that were marked as supplemental or secondary and/or where one or both mates did not map. BEDtools “bamtobed” (-bedpe; version v2.30.0) [73] was used to convert BAM records to “BEDPE” format. AWK was used to convert BEDPE format to Juicer’s “extra short” format, from which a HIC file (“.hic”) was created using JuicerTools (min. MAPQ of 10; v2.20) [69]. The HIC file was visualized in JuiceBox (v2.15) [67]. Note that the procedure described above for aligning, filtering, and visualizing Hi-C data was used to create all Hi-C map figures of Bcop_v1, Bcop_v1_corrected, and Bcop_v2 that appear in this manuscript.

JuiceBox [67] visualization aided manual inspection of the Hi-C signal across Bcop_v2 corresponding to individual chromosomes and individual contig joins within chromosomes. BED files for optical map gaps were used to visualize their locations and help interpret drops in Hi-C signal and depth of coverage on the Hi-C maps. BED files of Bcop_v1_corrected contig locations across Bcop_v2 were used to visualize them as squares across individual chromosome-scale scaffolds for manual inspection of the Hi-C-guided contig joins. To visualize and manually inspect contig joins in IGV [72], we extracted just the interaction frequencies between the start or end of a selected contig (left-most or right-most 1 kb or 10 kb of the contig) with 1 kb (or 10 kb) bins corresponding to the rest of the chromosome it resided on. Briefly, to do so we extracted “target-linked mates”, which are Hi-C paired-end reads that had at least one of the mates mapped within the targeted bin. Target-linked mate locations (interactions) in BEDPE format were visualized as arcs or links in IGV, which all emanate from the target region bin. However, although visualizing the links this way helps see all regions that are connected to the target bin by paired reads, it vastly misrepresents the interaction frequencies. Therefore, to visualize interaction frequencies across the scaffold, BEDtools was used to count the number of target-linked mates in bins that were constructed on the contig-bearing scaffold starting from the star/stop boundaries of the target region. Since interaction frequency decays with distance, log10 was also used (after adding a pseudocount of 1 to all bins) to better visualize long-distance interactions, such that a log10 score of 0 represented 0 interactions.

### Testing coverage expectations of the X chromosome compared to autosomes

Sequencing and mapping coverage across the four chromosome-scale scaffolds was visualized using a variety of orthogonal genomic datasets produced with different technologies and different life stages from previous studies [9,10]. In sum, we used Oxford Nanopore MinION and PacBio long reads from embryonic gDNA [9], BioNano Genomics long-range optical maps from male pupae gDNA [9], and Illumina paired-end reads from male and female adult gDNA [10]. Paired-end short reads were mapped as in Baird et al (2023) [10]. All long reads were mapped with Minimap2 with technology-specific settings [74]. The optical maps were aligned with Maligner [75] with the following considerations: (i) the Bcop_v2 reference genome sequence was converted to ‘*in silico* optical maps’ (make_insilico_map with CACGAG recognition sequence); (ii) the *in silico* Bcop_v2 maps and raw optical map data were “smoothed” using smooth_maps_file with “-m 1500” and “-m 600”, respectively; and (iii) smoothed optical maps were aligned to the smoothed *in silico* Bcop_v2 maps with maligner_dp (“--max-alignments 1” and otherwise default parameters). Genomic DNA sequencing coverage was computed for each dataset using BEDtools [73] and converted to bigWig using UCSC Kent Utilities [76] for visualization in IGV [72]. Coverage from long reads was computed after filtering for a minimum read length of 5 kb and a minimum MAPQ of 40. Coverage from BioNano optical maps was computed after filtering for molecules with a minimum length of 150 kb that aligned to Bcop_v2 with a maximum Maligner M-score of −10.

### Gene annotations for the chromosome-scale assembly (Bcop_v2)

To lift-over gene annotation sets originally produced on Bcop_v1, we used the program designed for this task, LiftOff [77] (version v1.6.3) with Minimap2 [74] (version 2.24-r1122). Along with the Bcop_v1 and Bcop_v2 assembly FASTA files and the Bcop_v1 GFF file, the following parameters were given to LiftOff: -p 16 -u unmapped.txt -polish -flank 0.25 -a 0.9 -s 0.9. The “polished” and “unpolished” outputs were identical for the Maker2 annotation, and were identical except for one gene for NCBI, which was extended 115 bp. Manual inspection showed the “unpolished” version to be correct in terms of matching Bcop_v1. Therefore, the “unpolished” versions are provided. GFF files for both gene annotation sets were deposited to: https://github.com/JohnUrban/Bcop_v2.

To compare the lifted-over gene models to RNA-seq evidence, RNA-seq datasets were first mapped genome-wide to Bcop_v2 without reference to any gene models using STAR [78] with the following parameters: --runThreadN ${P} --genomeDir ${STARIDX} --outSAMtype BAM SortedByCoordinate --readFilesCommand zcat --outFilterType BySJout --outFilterMultimapNmax 20 --alignSJoverhangMin 8 --alignSJDBoverhangMin 1 -- outFilterMismatchNmax 999 --outFilterMismatchNoverLmax 0.04 --alignIntronMin 20 -- alignIntronMax 1000000 --alignMatesGapMax 1000000 --outSAMattributes NH HI AS NM MD --sjdbScore 1 --readFilesIn ${R1} ${R2}. Resulting BAM files were indexed with SAMTools [66]. Genome-wide coverage was computed two ways for comparison: (i) with BEDtools [73] using the genomecov (-split -bg -ibam) and sortBed functions to produce a bedGraph that was converted to bigWig using bedGraphToBigWig from UCSC Kent Utilities [76]; and (ii) with RSEM [79] utilities using rsem-bam2wig (--no-fractional-weight) to produce a Wig file that was converted to bigWig using wigToBigWig [76]. The gene model GFF files produced by LiftOff [77] and the bigWigs for RNA-seq coverage profiles across Bcop_v2 were viewed in IGV [72]. The gene model examples shown for RNA-seq coverage are as follows, adorned with Maker2 (Bcop_v1) [9,34] and NCBI RefSeq (gene-LOC) [35] identifiers, either of which can be used with the lifted-over GFFs to find the coordinates in Bcop_v2. Bcop-PALP1: Bcop_v1_g007065; gene-LOC119085384. AGO1-like: Bcop_v1_g003309; gene-LOC119067568. Aub-like: Bcop_v1_g01302; gene-LOC119068172. CYP53A57: Bcop_v1_g020678; gene-LOC119075328. CYP3718B1: Bcop_v1_g007236; gene-LOC119074427. CYP4721A1: Bcop_v1_g009136; gene-LOC119078932.

To compare male and female expression of lifted-over gene models grouped by chromosome, we used published RNA-seq data for males and females across various life stages [9]. Transcript level quantification of all genes in the NCBI annotation set [35] lifted over to Bcop_v2 was carried out by RSEM (v1.3.1) [79] coupled with STAR (version 2.7.10a_alpha_220818) [78] using the “rsem-calculate-expression” pipeline. The RSEM-calculated expected counts were used with EdgeR [80] for between-sample normalization and differential expression analysis. Plots of EdgeR log2 fold-change results were made in Python.

### Using comparative genomics and evolutionary expectations to test the groupings of Bcop_v1 contigs into chromosome-scale scaffolds (Bcop_v2)

We identified orthologous relationships across all proteins within the proteomes of the five species using OrthoFinder [81,82], which also estimated the proteome-wide rooted species tree. For *B. coprophila*, we used NCBI *Bradysia coprophila* Annotation Release 100 protein set for Bcop_v1 (NCBI GenBank GCA_014529535.1; RefSeq GCF_014529535.1) [35] and the corresponding gene coordinates lifted over to Bcop_v2 described above. For the other species, we used the protein sets and genomic GTF files associated with the following NCBI accessions: *Drosophila melanogaster* (NCBI GenBank GCA_000001215.4; RefSeq GCF_000001215.4) [40]; *Pseudolycoriella hygida* (NCBI GenBank GCA_029228625.1) [42]; yellow fever mosquito (Aedes aegypti; NCBI GenBank GCA_002204515.1; RefSeq GCF_002204515.2) [41]; African malaria mosquito (Anopheles gambiae; NCBI GenBank GCA_000005575.1; RefSeq GCF_000005575.2) [39]. The proteome-wide rooted species tree was visualized using icy tree [83] by providing the Newick code returned by OrthoFinder [81,82]: “(D. mel:0.243236,((A. gam:0.26685,A. aeg:0.227034)0.954721:0.17008,(P. hyg:0.177818,B. cop:0.123002)0.969863:0.240218)1:0.243236);”.

OrthoFinder [81,82] creates distinct “Ortho Groups” containing all orthologs for a given gene from all given species. Thus, one can use the genes within an Ortho Group to compare the genomic locations of the “same” gene across all species. For simplicity and higher specificity, we used only the subset of Ortho Groups that are composed of a single ortholog from each and all of the five species, otherwise known as Single Copy Orthologs (SCOs). Thus, using the genomic location of a single gene from each species in an orthogroup of SCOs, we can make dot plots (or scatter plots), that show the location of the “same” gene in one species’ genome on the X-axis and another species’ genome on the Y-axis. Such dot plots were made in R using the assembalign.dotplot function from Lave [9,84] (https://github.com/JohnUrban/lave). Note that we used “conserved fly genes” in other analyses, which were *B. coprophila* genes that had one or more orthologs in all five Dipteran species, as opposed to only SCOs (which have one and only one copy in all five species).

We quantified the amount of “gene shuffling” using entropy, and implemented this analysis in Python3. Briefly, entropy can be thought of as a measure of uncertainty, information, disorder, or “mixed-up-ness”, and is often used to study systems that evolve from a highly ordered to a highly disordered state or from a non-random to a random state [44]. Mathematically, entropy is defined as −1 times the sum of p*log(p) for all probabilities, p, in a set of probabilities, P. That is, −1*sum(p*log(p)). When log base 2 is used, entropy is expressed in “bits”. For “gene shuffling”, the initial non-random (lowest entropy) state corresponds to how genes are grouped on chromosomes within a given species. As genes get shuffled around into different chromosomal groupings across evolutionary time, they approach randomized (high entropy) states with respect to some target species (e.g., *B. coprophila*). To calculate the entropy between any two species, we used only SCOs that appeared on the chromosome-scale scaffolds of each species, which is to say we excluded those that appeared on unplaced contigs. We counted the number of SCOs that appeared on each pair of inter-species chromosomes (i.e. chromosome “i” from species A and chromosome “j” from species B), added a pseudo-count of 0.1 to the SCO count of each inter-species pair, then divided each inter-species pair SCO count by the sum of SCO counts across all pairs of inter-species chromosomes (including pseudo-counts). Note that, for *D. melanogaster and A. gambiae,* the SCO counts from the left and right arms of chromosomes 2 and 3 (2L, 2R, 3L, 3R) were summed to get the full counts for those chromosomes (2L+2R, 3L+3R). For each pair of species, this gave a table of joint probabilities of a SCO appearing on chromosome “i” from species A and chromosome “j” from species B (see Table 6 for an example). The inter-species entropy was computed from these joint probabilities using the entropy equation: −1*sum(p*log2(p)), where p is the matrix of joint probabilities. However, as each pairwise species comparison has a different lowest possible entropy state (the non-random initial state) and a different highest possible (fully random) entropy state given their SCO counts per chromosome, we needed to use min-max normalization ((X-min)/(max-min)) in order to compare entropy values across pairwise inter-species comparisons. To calculate the minimum (initial) entropy state for each species within the context of a pairwise comparison, we marginalized (i.e. summed) over the joint probabilities to get the marginal probabilities that a SCO came from each chromosome for a given species. The entropy of the marginal probabilities of both species was then computed separately, using the equation above, and averaged together to get the lowest entropy value the joint probabilities could take. When comparing a species to itself, the average simply returns the entropy of the marginal probabilities for that species. To calculate the maximum (fully random) entropy state between two species, the marginal probabilities for each chromosome of a given species were multiplied by the marginal probabilities for each chromosome of the other species. This yielded a table of fully-random joint probabilities where the probability of a SCO appearing on chromosome “i” from species A and chromosome “j” from species B was simply the product of the marginal probability of a SCO appearing on chromosome “i” from species A and the marginal probability of a SCO appearing on chromosome “j” from species B. For the given chromosomal SCO counts for each species, this produces the joint probabilities expected at random, and maximum entropy is computed on it as above: −1*sum(p*log2(p)), where p is the matrix of random joint probabilities. The Seaborn library [85] in Python3 was used to visualize the min-max normalized entropy scores calculated against *B. coprophila* as a bar plot and all pairwise entropy scores as a clustered-heatmap.

### Testing gene locations of “alien” P450 genes within the chromosome-scale scaffolds using a variety of genomic datasets

If the “alien” cytochrome P450 (“CYP”) genes reported recently [7] are contained within the nuclear chromosomes, then they should exhibit expected Hi-C interaction frequencies [11] within the scaffolds. If they are external contamination erroneously stitched into the scaffold sequences, there should be severe drops in the Hi-C interaction frequency signal with these genes and the rest of the scaffold they reside in. To obtain all Hi-C interactions between a given “alien” P450 gene and all other sites in the genome, the same procedure described above for contig ends was used, here using Hi-C paired-end reads that had at least one of the mates mapped within the gene. For each “alien” P450 gene, the closest highly conserved fly gene that had a similar length (within 1% difference) was also interrogated the same way. Interactions and interaction frequencies in 100 kb bins were computed and visualized in IGV similar as described above, but with important differences. The number of interactions between a gene and the rest of the genome, and whether interactions are sparse or dense, is a function of gene length. To make valid comparisons with nearby conserved fly genes, gene length was controlled for in two ways. The first was to use nearby conserved fly genes of similar size (within 1%), as stated above. This helped match the observed interaction sparsity or density. The second was to normalize counts to each gene’s length, then multiply by 10,000 such that the magnitude of counts for all genes corresponded to a gene length of 10 kb, which was chosen since the “alien” gene lengths spanned from ∼1.8-7.8 kb.

Read depth can also differ significantly at sites of contamination, especially across different biological samples. Some samples may show low or no sequencing depth over sites where contamination was integrated into a genome assembly. Other samples with an abundance of the contaminant may show spikes in read depth there. It is unlikely for such sites of integrated contamination to have no disruptions in the depth signal across multiple samples. To investigate the DNA-sequencing coverage over the regions bearing “alien” P450 genes, we queried the sequencing depth (computed as above) of a diverse set of biological samples, including different sexes and developmental stages [9,10], using datasets from orthogonal technologies, including Illumina-paired end reads [10], confidently-mapped (MAPQ≥40) long (≥5 kb) Nanopore and PacBio reads [9], and stringently-mapped ultra-long (≥150 kb) optical maps from BioNano Genomics [9]. See above “coverage” Methods subsection for details.

### Repeat annotations for the chromosome-scale assembly (Bcop_v2)

RepeatMasker [71] was used to re-annotate the repeats across Bcop_v2 using comprehensive repeat libraries as done previously for Bcop_v1 [9], rather than using a lift-over process. As previously, the comprehensive repeat library contained (i) species-specific repeat libraries learned with RepeatModeler [86], (ii) previously known repeat sequences from *Bradysia coprophila* found at NCBI prior to Bcop_v1 [9], and (iii) all Arthropod repeats in the RepeatMasker Combined Database, which included Dfam_Consensus-2018/10/26 [87] and RepBase-2018/10/26 [88]. The corresponding repeat annotation GFF file for Bcop_v2 was deposited to: https://github.com/JohnUrban/Bcop_v2. RepeatMasker [71] was also used to map just the arthropod repeats alone. TandemRepeatFinder (TRF) [89] was used independently to map and visualize just tandem repeats in the genome (trf ${FA} 2 7 7 80 10 50 2000 -f -d -m). Repeat densities in 100 kb bins across the genome were computed using BEDtools [73]. Briefly, either all repeats in the RepeatMasker GFF or targeted repeats (e.g. a given family like LINEs) were extracted and converted to BED; the proportion of bases corresponding to those repeat intervals within 100 kb windows was calculated using the “coverage” BEDtool [73]. Repeat density bedGraphs were visualized in IGV [72]. Retrotransposon-related sequences and satellite sequences were found by inspecting the repeat annotation and tandem repeat finder results in the first 10 kb (or up to 100 kb) of the scaffold termini using IGV [72], and looking at dot plots of the first 10-100 kb. Their distributions across the scaffolds were further explored by BLASTing [70] them back to the assembly.

Chromosome-wide and genome-wide dot plots were made to visualize the repeat structure across the scaffolds. Briefly, Minimap2 [74] was used to map chromosomes against themselves or the entire genome against itself using parameters optimized for dot plots (-PD -k19 -w19 -m200 - t8). The PAF output files were visualized as dot plots in R using the assembalign.dotplot function from Lave [9,84], which presents the pairwise-aligned query and target intervals as line segments. BED files for the locations of anchor sequence midpoints +/- 500 kb, estimated centromere midpoints +/- 500 kb, and estimated locations of Fold-Back Regions 1-3 and the X’ long paracentric inversion breakpoints were provided to annotate the plot margins for reference points.

### Extracting and analyzing locus sequences corresponding to long-range Hi-C interactions on the X chromosome

The locations of the loci on Fold-Back Regions on the X chromosome that appear as three off-diagonal dots corresponding to long-range interacting loci on the Hi-C maps, as well as the “main breakpoints” corresponding to a long paracentric inversion with the X’ chromosome variant, were learned by analyzing the paired Hi-C reads that mapped to the X with long distances in between them. Two datasets were explored: male (XO) pupae data from this study as well as female X’X Hi-C data from a recent analysis of the X’ chromosome [10]. Briefly, paired-end Hi-C reads (in Juicer’s Extra Short format [69]) that mapped to the X chromosome using the above pipeline were read into R where map locations were further processed to filter for paired reads with (i) inter-mate distances of at least 1 Mb, and (ii) minimum mapping qualities (MAPQ) of 10, although a range of MAPQs were explored, including no filtering, and gave similar end results. The 2D Binned Kernel Density Estimate (from the ‘KernSmooth’ package) was then taken using a 1024×1024 grid (∼68.9 kb bins) with 250 kb smoothing bandwidths on both the x- and y-axes corresponding to where mate1 and mate2 mapped across the X. During optimization and exploratory analyses, kernel-smoothed 2D grids were visualized using levelplot from the ‘lattice’ package. Peaks in the Z (heat) dimension of the 2D grid were called by identifying x,y-coordinates where the Z dimension was above the 99.8th percentile across the entire grid. In other words, Z-peaks corresponded to pairs of distant loci (>1 Mb) with the highest numbers of Hi-C reads connecting them. The x,y-coordinates of peaks in the 2D grid were then hierarchically clustered, and the tree was cut to identify k=12 peak clusters on the 2D grid. The median, minimum, and maximum x- and y-coordinates of each cluster were obtained to help define the centers and the boundaries of interacting pairs of loci on the X chromosome, as was the coordinates of the peak summit bin (the bin with the highest interaction frequency in a peak cluster). The peak clusters were filtered for those whose median x-coordinates were separated by 5 Mb from the median y-coordinates. In other words, these were loci pairs separated by at least 5 Mb. Three of the four remaining peak clusters separated by at least 5 Mb corresponded to the interactions between putative FBRs 1 and 2, putative FBRs 1 and 3, and putative FBRs 2 and 3. The fourth remaining peak cluster corresponded to the breakpoint dots for the long paracentric inversion with the X’. The peak clusters for loci separated by less than 5 Mb may correspond to (i) other smaller structural variations between the X’ and X, and (ii) other shorter Mb-range interactions of distal loci. The regions corresponding to the three FBRs and the two main X’ breakpoints were visualized on Hi-C maps using JuiceBox [67]. Note that using inter-mate distances of 5 Mb instead of 1 Mb and k=4 clusters gave the same end results without needing further filtering. With these parameters peak clusters ranged from 551 kb to 1.1 Mb (8-16 bins) and had summit bins of 68.9 kb. We also identified parameters for much narrower estimates of FBR locations using MAPQ of 10, intermate distance minimum of 5 Mb, a 2048×2048 grid (∼34.4 kb bins) with a smoothing bandwidth of 50 kb, a peak quantile threshold of 0.9999, and k=4 peak clusters, which produced four interaction peak clusters in the ∼103-551 kb range (3-16 bins) with summit bin estimates of ∼34.4 kb.

The nucleotide distances between the mid-points of the FBR regions, X’ breakpoints, and centromeres observed on the X scaffold were compared to expected nucleotide distances given corresponding polytene map distances in proportion to the length of the X chromosome scaffold. The X polytene map is organized into 14 zones of similar lengths, each with 3 subzones (A, B, and C) for a total of 42 sub-zones. Assuming nucleotide lengths of zones are also similar, the ∼70.5 Mb X-scaffold would indicate zone lengths of ∼5 Mb and sub-zone lengths of ∼1.68 Mb, which we used to compute expected lengths. The FBR coordinates were used to define both the sequences of the FBRs as well as the Genomic Complement (the genome sequence excluding the FBRs). FBR and Genomic Complement sequences were extracted using BEDtools [73]. JellyFish [90] was used for kmer operations on both sets of sequences. BLAST [70] was used to compare FBR sequences to each other and to the Genomic Complement sequences. FBR dotplots were made as described for the entire genome and individual chromosomes above. BEDtools [73] was used to extract and analyze genes and repeats in the FBRs. The FBR coordinates were also used to annotate the X chromosome dotplot, IGV traces showing repeats, and Hi-C interaction frequency maps.

## Data Availability

This Whole Genome Shotgun project has been deposited at DDBJ/ENA/GenBank under the accession VSDI00000000. The version described in this paper, referred to throughout as Bcop_v2, is version VSDI02000000 (deposited 2022; released Jan 4, 2023). This project is associated with BioProjects PRJNA291918 and PRJNA672144; and BioSamples SAMN12533751 and SAMN20343824. Male Pupa Hi-C data was deposited to SRA: SRR23335771. Lifted over gene model GFFs, repeat annotation GFF, *B. coprophila*-specific repeat libraries, as well as AGP and BED files mapping Bcop_v1 to Bcop_v2 were deposited to a github repository: https://github.com/JohnUrban/Bcop_v2. The RNA-seq samples from *B. coprophila* from different life stages used to analyze dosage compensation and test gene models are from Urban et al (2021) [9] and associated with BioProject PRJNA748150. The RNA-seq samples from the radiation study used to test gene models and visualize exon coverage are from Urban et al (2023) [37] and associated with BioProject PRJNA928089. The long reads from Oxford Nanopore and PacBio, and optical maps from BioNano Genomics are from Urban et al (2021) [9] and associated with BioProject PRJNA291918, and the Illumina reads from male and female adults as well as X’X Hi-C data are from Baird et al (2023) [10] and associated with BioProject PRJNA953429.

## Abbreviations

AGP: a golden path (file format)

Bcop: *Bradysia coprophila*

bp: base pairs

BED: Browser Extensible Data (file format)

BUSCO: Benchmarking Universal Single-Copy Orthologs

CDS: coding sequence

FBR: foldback region

Gb: gigabase pairs

GFF: General Feature Format (file format)

GTF: General Transfer Format (file format)

Hi-C: High dimensional Chromosome conformation capture

i5k: initiative to sequence 5000 insect (and other arthropod) genomes (http://i5k.github.io/about)

kb: kilobase pairs

LCTTR: long complex terminal tandem repeats

LINE: long interspersed nuclear element

LTR: long terminal repeats

Mb: megabase pairs

MMNE: Min-Max Normalized Entropy

NCBI: National Center for Biotechnology Information

PacBio: Pacific Biosciences

PCR: polymerase chain reaction

RTE: retrotransposable element

RTRS: retrotransposon related sequence

Sccr: *Sciara coprophila* centromeric repeat

ScRTE: *Sciara coprophila* retrotransposable element as defined in Escribá et al (2011) [45]

SMRT: single-molecule real-time.

ONT: Oxford Nanopore Technologies.

## Authors’ Contributions

JMU, SAG, and ACS conceived the project. SAG collected male pupae, coordinated with Phase Genomics, and helped with chromosome maps and map positions. Hi-C library preparation, sequencing, and scaffolding was performed as a service provided by Phase Genomics. JMU performed the original Bcop_v1 assembly, and corresponded with Phase Genomics during the optimization process for Bcop_v2 scaffolding. JMU performed all subsequent analyses and operations, including subsequent Hi-C data processing, analysis, exploration and visualizations in all figures, chromosome identification, scaffold orientation, assembly evaluations, gene annotation lift-overs, comparative genomics and entropy analyses, HGT gene validations, repeat annotation and analyses, scaffold termini characterization, and fold-back repeat sequence extraction and characterization; JMU developed the ideas, analyses, and python/R code for Lave dot plots, for entropy/MMNE analyses, and for the long-range Hi-C interaction peak detection method to discover FBR and breakpoint locations; JMU made all figures and tables. ACS helped conceive of analyses, figures, and tables. All authors participated intellectually in the development and execution of this project. All authors wrote, read, and edited the manuscript.

## Acknowledgements

We thank Jacob Bliss who helped with *Bradysia coprophila* maintenance during data collection. We thank our correspondents at Phase Genomics, including Hayley Mangelson, Kayla Young, Shawn Sullivan, Brian Fan, Andrew Wiser, and Ivan Liachko. We acknowledge and thank Rob Baird and Laura Ross for being early adopters of Bcop_v2 for collaborative studies of the X’ chromosome. We thank Jennifer Urban for proof-reading the manuscript. We thank Ava Filiss for the X and X’ polytene chromosome images. We thank all members of the Spradling Laboratory for feedback during lab meetings, and especially Robert Levis for discussions on telomeres. Polytene chromosome maps in figures 3, 7, and 8 were reproduced from plates 1, 2 and 3 of Gabrusewycz-Garcia (1964) [30] with permission from Springer Nature under permission number: 5490830617309. We thank bioRxiv for hosting our 2022 preprint that preceded this paper (https://doi.org/10.1101/2022.11.03.515061; version 1, Nov 2022; version 2, Dec 2024).

